# Dairy cattle movement networks govern the spread and persistence of H5N1 B3.13 in the United States

**DOI:** 10.64898/2026.10.28.762465

**Authors:** Jonathan E. Pekar, Yucai Shao, Alvin Crespo-Bellido, Prathyush Sambaturu, Guy Baele, Joel O. Wertheim, Praneeth Gangavarapu, Karthik Gangavarapu, Gytis Dudas, Thomas P. Peacock, Alexander Maksiaev, Tanin Rajamand, Michael W. Sanderson, Justin Bahl, Ethan Plunkett, Adriaan M. Dokter, Jennine Ochoa, Kristian G. Andersen, Oliver Pybus, Michael Worobey, Martha I. Nelson, Moritz U. G. Kraemer, Marc A. Suchard, Andrew Rambaut, Philippe Lemey

## Abstract

Avian influenza virus H5N1 is predominantly found in wild and domestic birds. H5N1 genotype B3.13 was first detected in United States dairy cattle in March 2024 and spread through much of the country during 2024 and 2025, resulting in the largest known H5N1 outbreak in domestic mammals. By analysing viral genomic data jointly with historical cattle movement patterns, we find that live cattle movement is the predominant contributor to H5N1 spatial spread, rather than wild or domestic birds. Using phylogeographic methods we estimate that >40% of interstate viral transmissions occurred before B3.13’s first detection and >70% occurred before interstate dairy cattle movement restrictions were implemented in April 2024. After this implementation, interstate transmission slowed but within-state spread persisted, driven by local cattle network density and movement volume. Our epidemic simulations demonstrate that the connectivity of counties within and around the Texas Panhandle—where the earliest cases were documented—predisposed the region to become an outbreak epicentre. Counterfactual simulations indicate that interstate restrictions needed to have begun 2–3 months earlier in order to limit the epizootic substantively, given the extent of cryptic spread before detection. Network-informed interventions outperformed geographically-defined containment, but response speed (rather than intervention strength) was the limiting factor in our epidemic models, underscoring the need for surveillance systems capable of detecting pathogen emergence before widespread dissemination.

## Introduction

In early 2024, the avian influenza virus (AIV) H5N1 genotype B3.13 was identified in dairy cattle in the southern United States (US) and subsequently spread extensively among dairy-producing regions in the country^1^. The first outbreak was confirmed on 25 March 2024, and by late April, B3.13 had been detected on dozens of dairy cattle premises across 9 states. A federal order requiring pre-movement testing for interstate dairy cattle transport was implemented on 29 April 2024 (ref.^2^), but additional outbreaks continued to be detected, and by the end of 2025, there were 1,068 documented outbreaks across 17 states^3^.

The emergence of B3.13 in dairy cattle was likely due to a single spillover of B3.13 from wild birds in late 2023^1,4^. Since then, reports of dead peridomestic birds near dairy farms have raised speculation that repeated spillover could have contributed to B3.13’s geographic spread^5^, thereby bypassing cattle movement restrictions. The B3.13 lineage in dairy cattle is monophyletic, indicating its emergence in dairy cattle was due to a single introduction, but it is plausible that spillback into wild birds followed by reintroduction into cattle could retain monophyly while continuing to spread.

The establishment of B3.13 in the US occurred in the context of a heterogeneous landscape of industrial dairy cattle practices^6,7^. Interstate transmission among dairy cattle has been suggested to have seeded many outbreaks^1,5^, but regional-level patterns and drivers of the epidemic have remained uncertain for both within- and between-state transmission. Understanding the predictors of viral transmission across geographic scales, and which movement restrictions most effectively mitigate spread, is critical for responding to future livestock outbreaks.

Here we integrate pathogen genomic data and modelling approaches to analyse the spread of B3.13 across the US. We first reconstruct viral spread among wild birds, domestic birds, and dairy cattle, to determine whether transmission is predominantly sustained within dairy cattle populations. We then reconstruct a detailed phylodynamic history of B3.13 among dairy cattle in the affected states and test which state-level features are associated with the spatial dynamics of B3.13. Our analysis investigates B3.13 dissemination before its first confirmed detection and before and after subsequent implementation of movement restrictions. We further use epidemic modelling to examine which county-level features predict the emergence and multi-state spread of B3.13 across the dairy cattle movement network. Finally, we investigate how transmission dynamics varied by intervention scenarios, including restricting infectious herd movements and whether earlier or more stringent movement restrictions could have mitigated the epidemic. Our results elucidate how dairy cattle movement networks shape viral establishment, persistence, and dissemination across geographic scales.

## Results

### Wild birds did not contribute to long-range B3.13 dispersal

To determine whether sustained transmission of B3.13 among dairy cattle was predominantly due to cattle-to-cattle transmission, rather than recurrent cross-species transmission between cattle and wild birds, we performed a phylodynamic reconstruction employing an asymmetric discrete phylogeographic model^8^ using 1,358 genomes comprising three host groups: dairy cattle, domestic birds (poultry and captive birds), and wild birds. We inferred dairy cattle as the ancestral host of most internal nodes and branches, whereas wild-bird viruses occurred as singletons or as small terminal clusters nested within cattle-associated lineages (Figs. 1a, S1). The transition rate from dairy cattle to domestic birds was the only relative rate predominantly above 1 (Fig. 1b). Our analysis indicates that there has been spillover of B3.13 from dairy cattle into wild birds (median number of transition events=15, 95% highest posterior density [HPD]: 12–17), with a smaller number from domestic birds into wild birds (median=2, 95% HPD: 1–4). In contrast, we find minimal support for host switches of B3.13 from wild birds into either dairy cattle (median=0, 95% HPD: 0–1) or domestic birds (median=0, 95% HPD: 0–0; Fig. 1b).

**Figure 1.**
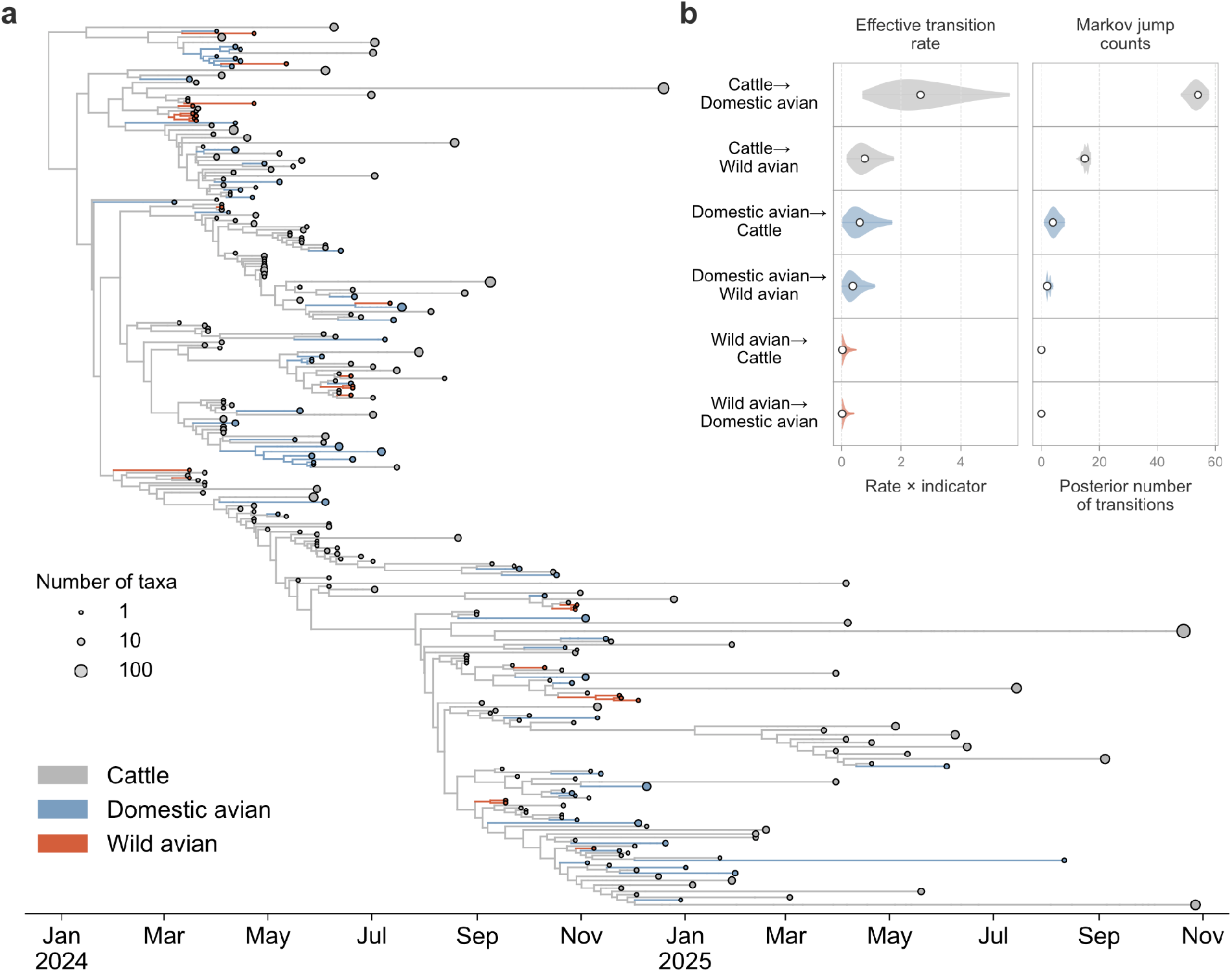
Multi-host transmission dynamics of B3.13. **a**, Time-calibrated phylogenetic tree of B3.13, with branches coloured by their most probable host. Monophyletic clades sharing a single host are collapsed into circles proportional to the number of constituent taxa, positioned at the location of the most recently sampled tip. Individual tips are shown where clades remain uncollapsed. Branch and circle colors indicate host identity. **b**, The effective host transition rate (left) and the posterior number of transitions between hosts (right). Violins are colored by source host and depict 95% highest posterior density (HPD), and white circles represent the median value. Figure S1 provides a summary of the observed spillover events.

To evaluate whether these results were biased by the relatively high number of dairy cattle virus genomes, we iteratively downsampled the number of dairy cattle virus genomes and repeated our phylogeographic inference. We continued to find limited support for transitions of B3.13 from wild birds to either dairy cattle or domestic birds (Fig. S2); only after downsampling dairy cattle to 12.5% of the original count, whilst keeping the number of wild bird genomes constant, did we find a slightly elevated number of host switches from wild birds to dairy cattle (median: 3, 95% HPD: 0–6). Our findings are therefore robust to variation in the representation of dairy cattle viruses; the additional transitions from wild birds into dairy cattle inferred under this high level of downsampling likely reflect the reduced representation of cattle-to-cattle transmission.

When inferring the ancestral locations of these viruses in our largest dataset (n*=*1,358), we found that all clusters of wild bird viruses were from the same state; there were no branches indicative of wild bird-associated transmission across state lines (Fig. S3). This finding, coupled with the absence of phylogenetic evidence of prolonged unsampled transmission in wild birds and likely undersampling of B3.13 in dairy cattle, suggests that wild birds are predominantly a dead-end host and not responsible for long-range geographic spread.

### Dairy industry and movement data predict B3.13 spread

To identify the factors that contribute to the spatial spread of B3.13 among dairy cattle, we analysed 3,950 H5N1 B3.13 genomes sampled from dairy cattle in 2024 and 2025, and constructed a Bayesian time-measured phylogeographic model that integrated industrial, epidemiological, and mobility data^9^. Candidate predictors of spatial spread included measures of dairy industry structure (state-level inventory, herd counts, milk productivity, and replacement rates)^10^, geographic distance, and two independent characterisations of interstate cattle movement based on Certificate of Veterinary Inspections (CVI) records (Table S1). One characterisation consists of daily interstate and intrastate cattle movement volumes, generated by USAMMv3 (ref.^11^); specifically, a fitted stochastic network model parameterised from 2009 CVI records (referred to here as ‘movement data’; Fig. S4). The other characterisation consists of yearly sums of interstate confirmed and inferred shipment volumes, based on Interstate Certificate of Veterinary Inspections (ICVI) records aggregated from April 2015 through March 2016 (see Methods; referred to here as ‘regulatory shipments’; Table S2).

Analyses using time-homogeneous and time-inhomogeneous models offered strong support (Bayes Factor [BF]>50) for movement data, milk production at origin, and replacement rates at origin as predictors of spatial lineage spread (Fig. S5); there was little evidence for regulatory shipment records as a predictor, despite these data spanning a more recent time period than the movement data. We therefore focused on the three supported correlates in the subsequent state-level phylogeographic analyses.

The first detection of an outbreak was on 25 March 2024, and a federal order requiring pre-movement testing for interstate dairy cattle transport was implemented on 29 April 2024. The detection itself may have altered producer behaviour independently of formal policy (*e.g.*, voluntary reductions in cattle movement or equipment sharing^12^, or tightened state-level entry requirements, as enacted by California in early April 2024^13^), so we defined three eras—before detection (“pre-detection”), between detection and federal order (“pre-response”), and after the order was enacted (“post-response”)—and allowed both the strength of the predictor effects and overall rate of spatial spread to vary across these three eras. When conditioned on the three supported correlates, we found that the movement data were positively associated with interstate viral lineage movement across all three eras. Milk production at the origin location was, on average, positively associated with interstate viral lineage movement during the pre-detection era, but this relationship was not consistent, and the dairy cattle replacement rate at origin during the post-response era was negatively associated with interstate viral lineage movement. The same patterns were observed when allowing for variable selection during the time homogeneous analysis (BF>10 for the aforementioned predictors during their relevant eras; Fig. S6).

We used our probabilistic model of spatial spread, informed by genomic, industrial, and movement data, to characterize the dynamics of spread of B3.13 among states. Although genomic and epidemiological data suggest epidemic incidence peaked in the second half of 2024 (Fig. 2a,b), we find that most interstate lineage movements occurred between February and April of 2024, before the enactment of the federal order, and that these movements peaked at nearly the same time as the first confirmed outbreak (median date = 20 March 2024; 95% HPD: 14 March–25 March; Fig. S7). The majority of viral lineage movements during the pre-detection era originated in Texas (Fig. 2c; n=19/29), but Michigan was a secondary and primary exporter during the pre-detection (n=7/29) and pre-response eras (n=8/21), respectively. Texas had few viral lineage imports relative to exports (Figs. 2d, S8, S9), and several states, such as Oklahoma, appeared to be viral sinks that did not export B3.13 to other states. The majority of interstate lineage transitions occurred during the pre-detection and pre-response eras, with a median of 43.8% of transitions (95% HPD: 36.8–50.8%) occurring before confirmation of the first outbreak and 24.6% (95% HPD: 18.4–31.4%) occurring after the implementation of the federal order. Only 8.5% (95% HPD: 5.6–11.1%) of transitions occurred >2 months after the order, suggesting a brief lag before its full effect. Era-specific parameters indicated that the overall rate of interstate viral lineage movement was higher in the pre-detection and pre-response eras than in the post-response era (Fig. S6c,d), consistent with the majority of interstate transitions occurring before the federal order. The frequency of weekly interstate viral transitions then decreased rapidly during the pre-response era (Figs. 2c,d, S7), suggesting that the identification of the outbreak in dairy cattle could have led to behavioural shifts independent of the federal order.

**Figure 2.**
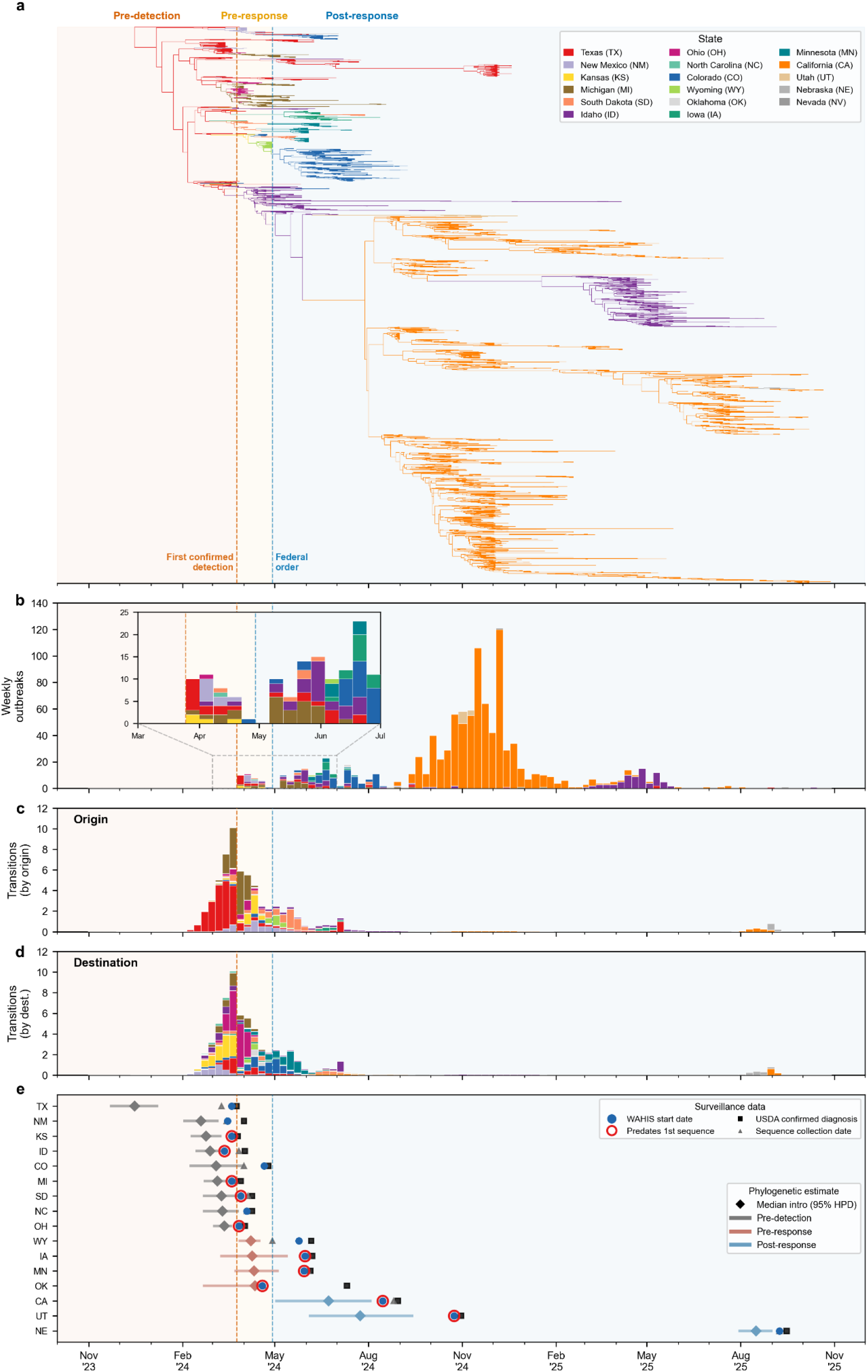
Emergence and spread of B3.13 in dairy cattle. **a**, Time-calibrated phylogenetic tree of B3.13 in dairy cattle, with branches colored by their most probable location (US state). Key shows US states with abbreviations in parentheses. **b**, Weekly outbreaks of B3.13 among dairy premises in the US. **c-d**, Posterior mean dates of inferred among-state B3.13 lineage transitions, colored by origin (c) or destination (d). **e**, Inferred date of introduction of B3.13 into each state, with posterior median indicated by a diamond and 95% highest posterior density [HPD] interval indicated by the solid line, colored by whether the median date is in the pre-detection era (grey; before the confirmation of the earliest B3.13 outbreak on a dairy premise, in Texas, 25 March 2024), the pre-response era (red; after the confirmation of the earliest outbreak but before the implementation of the federal order, 29 April 2024), or post-response era (after the implementation of the federal order). For Texas, the time of the most recent common ancestor (tMRCA) of the phylogeny is indicated, as Texas is the likely epicentre of the outbreak. The first reporting of the outbreak by WAHIS in each state indicated by a blue circle. The earliest USDA confirmation of an outbreak in each state indicated by a black square (Table S3). Gray trianangles show the dates of collection of the earliest genome from each state. If the date of the WAHIS reporting occurs prior to the collection of the earliest sequence, the blue circle representing the WAHIS reporting date is surrounded by a red circle. In each panel, the date of the earliest confirmed B3.13 outbreak is indicated by a dashed vertical orange line, and the date of the federal order implementation is indicated by a dashed vertical blue line. No B3.13 genomes are available from Nevada; it only appears in panel B.

We additionally observe notable surveillance gaps^14^—the lag between the introduction of the pathogen into a state and its detection—among the states with documented B3.13 outbreaks (Fig. 2e). We infer that introductions into states with confirmed outbreaks actually occurred at least 3–4 weeks before detection within that jurisdiction. Further, for four of the seven states with outbreaks first confirmed in the post-response era, the introductions of B3.13 occurred during the pre-response era and with even longer intervals between introduction and confirmation than observed for most states with confirmations in the pre-response era. The surveillance gaps for several states (*e.g.*, California and Utah) were likely two months or more, indicating substantial room for improvement in identifying and responding to detected outbreaks.

### Dairy cattle movement drives within-state persistence

The B3.13 outbreak continued at least until the end of 2025, but with limited interstate viral movements after the implementation of the federal order, indicating that within-state dynamics were critical in sustaining transmission. Virus persistence dynamics varied among US states, with some states sustaining few but extended transmission clusters (e.g., California; Fig. 3a) and others receiving more introductions that each persisted only briefly (e.g., Kansas). Variance decomposition of log-transformed total branch time confirmed this pattern: differences among states reflected how long each introduction lasted (85% of variance) more than the number of introductions (27%), suggesting state-level structural features were responsible for viral persistence.

**Figure 3.**
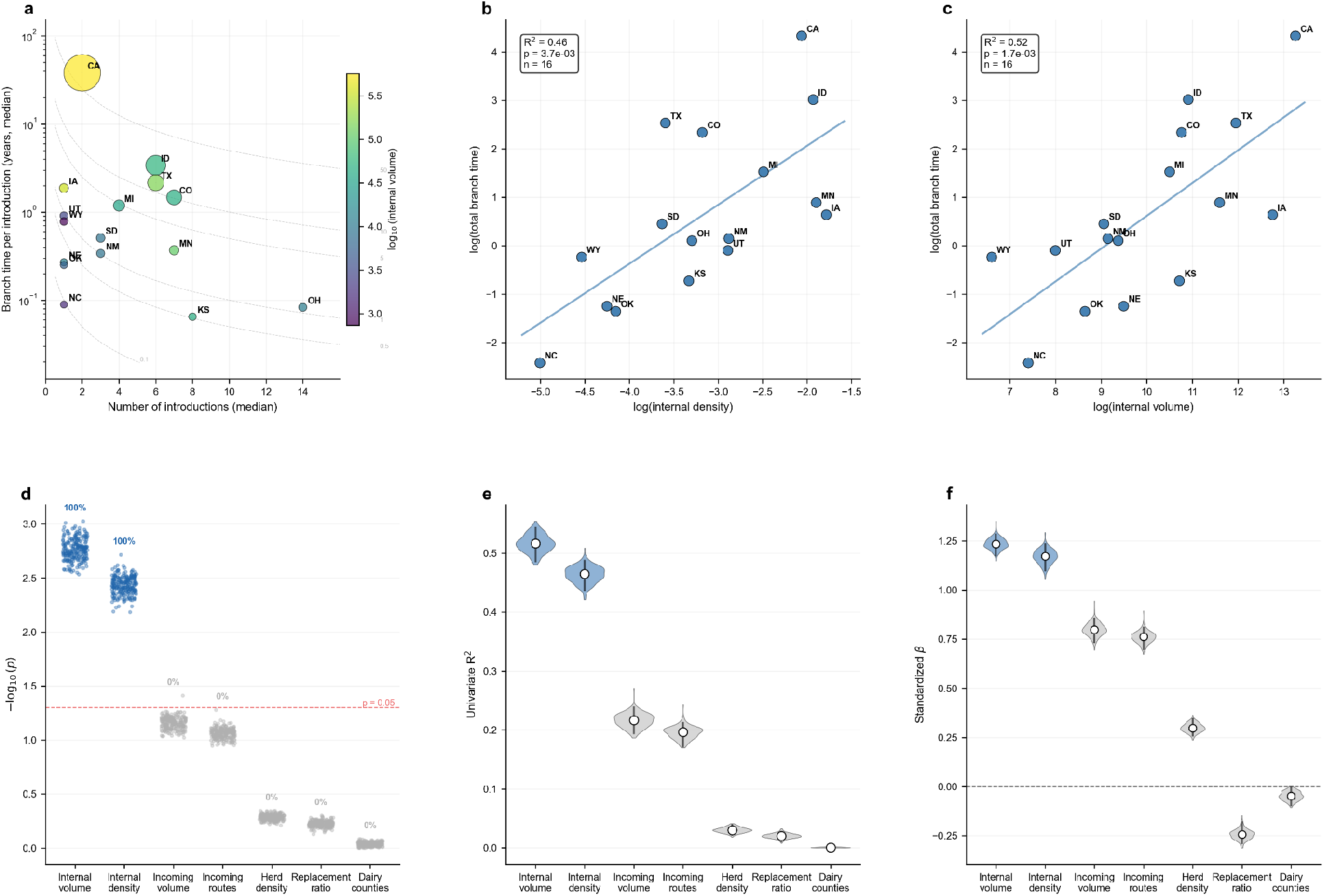
Within-state persistence of B3.13 is driven by internal cattle movement. **a**, Decomposition of total branch time into number of introductions and time per introduction for each state. Bubble size proportional to posterior median total branch time; color indicates log(internal cattle movement volume). Dashed isolines indicate equal total branch time. **b**, Relationship between log(internal network density) and log(total branch time). **c**, Relationship between log(internal cattle movement volume) and log(total branch time). R² and P-values are shown for regressions in b–c. **d**, Values of -log(P) for each predictor across the posterior distribution; each point represents one posterior tree sample. Red dashed line indicates the significance threshold (P = 0.05), with points above the line denoting statistical significance. Percentages indicate the proportion of posterior tree samples with P < 0.05. **e**, Univariate R² for each of the seven network and dairy predictors (bars, 95% HPD). **f**, Standardized regression coefficients β_i_ for each predictor. White dots indicate medians and thick bars indicate 95% HPDs. Blue violins denote predictors supported in ≥90% of posterior trees; grey violins denote predictors below this threshold. Dashed line indicates β = 0.

The movement data we used to parameterize our phylogeographic model also capture within-state movement structure, enabling us to interrogate which variables drive lineage persistence, including shipping routes, movement volume, and network density (Table S4). We screened each candidate variable independently against total within-state branch time (Fig. S10) and found that two predictors were consistently significant: internal network density (R^2^=0.46, *p*=0.0037; significant in 100% of posterior trees; Fig. 3b) and internal cattle movement volume (R^2^=0.52, *p*=0.0017; significant in 100% of posterior trees; Fig. 3c). The remaining five variables showed no consistent association (Fig. 3d,e). When screening each candidate independently against average branch time per introduction (Fig.

S10), our results were broadly consistent (Fig. S12). Sensitivity analyses consistently indicated support for the network density and cattle movement volume predictors (Fig. S13). However, these two variables are correlated (r = 0.70; Fig. S11) and cannot be reliably separated here, hence we interpret them as reflecting the same underlying driver: US states with more active internal cattle movement networks likely sustained longer chains of transmission following introduction, consistent with a greater degree of contact between the susceptible and infected populations^15,16^.

### Defining an outbreak epicentre

Given the established role of network connectivity in shaping epidemic dynamics, we sought to examine whether the connectivity of a putative county of origin can predict successful multi-state spread (here defined as infections that result in spread to at least two other states). We first constructed a parameterised framework that recapitulates the observed epizootic (Supplementary Note 1; Figs. S14–S18). Using this framework, we then simulated epidemics seeded from counties across the 16 states with documented B3.13 outbreaks, accounting for the federal order that subsequently reduced interstate transmission probability. Multi-state spread was strongly correlated with the mean out-degree of the seed counties in the top quartile of connectivity within each state (Fig. 4a), while bottom-quartile counties in the same states were highly unlikely to produce sustained outbreaks (Fig. 4b).

**Figure 4.**
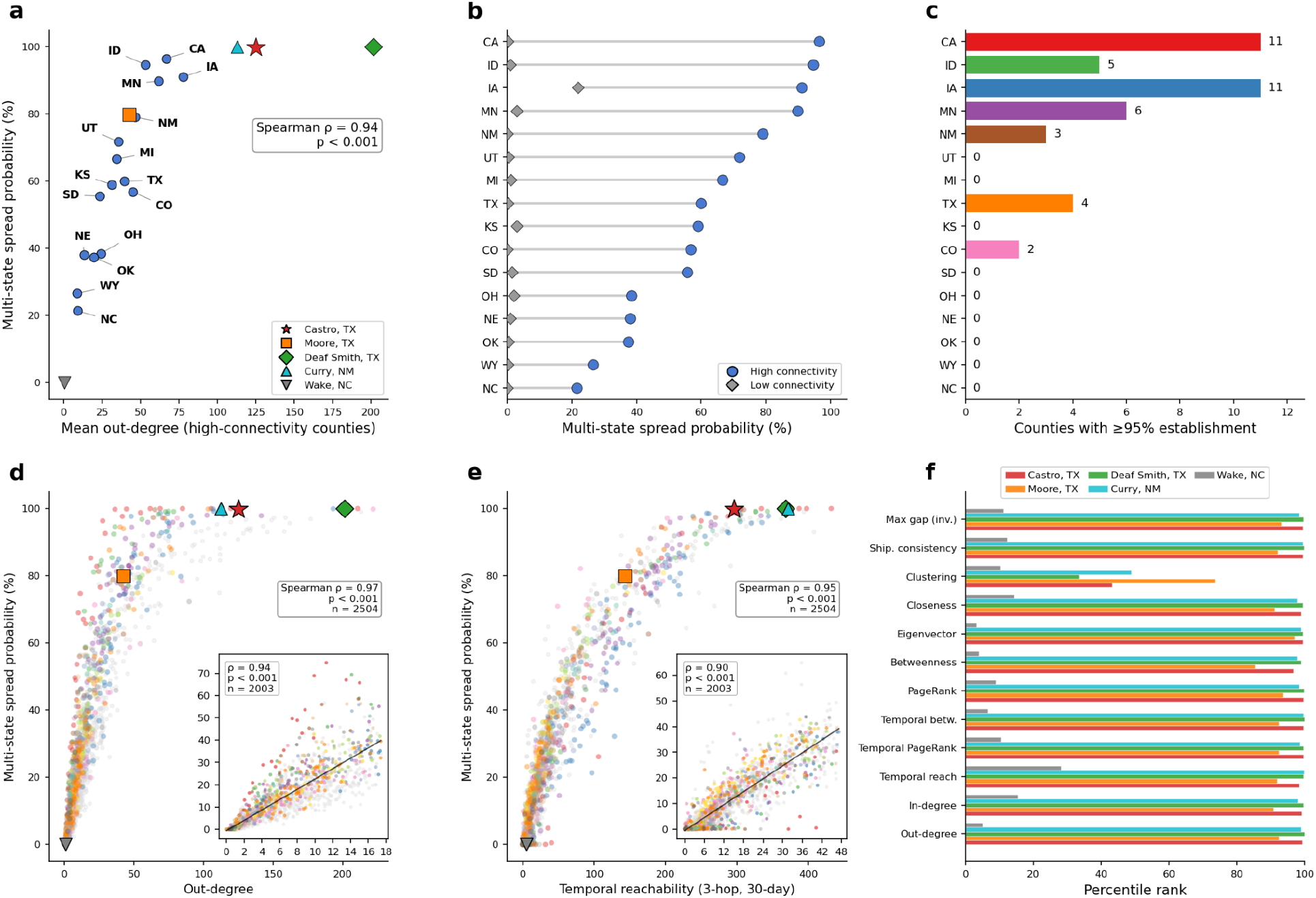
Network properties associated with viral spread. **a**, State-level correlation between mean out-degree and multi-state spread probability, computed from high-connectivity seed counties across the 16 states with observed outbreaks. **b**, Average multi-state spread probabilities of low-connectivity and high-connectivity seed counties across the 16 states with observed outbreaks. **c**, Number of counties with ≥95% multi-state spread probability across the 16 states with observed outbreaks. **d**, Correlation of multi-state spread probability with county out-degree across all 2,504 dairy counties in the 48 contiguous states, with inset showing counties below the 80th percentile of out-degree. **e**, Correlation of multi-state spread probability with temporal reachability, with inset showing counties below the 80th percentile of temporal reachability. **f**, Network percentile rankings for five counties from which many of the earliest reported cases were documented. These counties are also indicated in panels a, d, and e. Multi-state spread probability is defined as the probability of spreading to at least two additional states besides the one where the simulated outbreak started. Figure S20 shows multi-state spread probability across all 48 contiguous states.

The four counties with the earliest sampled genomes (Moore, Castro, and Deaf Smith Counties in Texas, and Curry County in New Mexico) all have high mean out-degree and multi-state spread probability (Fig. 4a). Conversely, Wake County in North Carolina, the source of a single early genome and the only inferred importation into the state, has minimal connectivity and near-zero multi-state spread probability. Multiple states beyond Texas contained counties where multi-state spread is near-certain, including California and Iowa (Figs. 4c, S20).

In addition to connectivity, the movement data (USAMMv3) allowed us to examine a range of structural and temporal network features (Fig. S19, Table S6). Extending our simulations to account for all counties with dairy cattle across the 48 contiguous states, we find that county out-degree (the number of edges out of a county; Fig. 4d) and temporal reachability (the number of counties reachable within 3 hops and 30 days across network realizations; Fig. 4e) were strongly correlated with multi-state spread probability. To confirm this finding was not driven by counties with extreme values for either metric, we repeated the analysis after excluding the top quintile and found that both metrics are still strongly correlated with multi-state spread probability (Fig. 4d,e, inset panels). The four counties with the earliest sampled genomes rank consistently near the top across the full array of network features (Figs. 4f, S19).

When seeded from high-connectivity counties, simulated outbreaks often reached over 30 states and over 400 counties, compared to a median of 1 state across all dairy counties (Fig. S21). Collectively, these results indicate which counties were likely to sustain onward transmission and reinforce why the Texas Panhandle and the immediately surrounding region, which together contain many such counties, was the outbreak epicentre.

### Mitigation and predicted impact of early control strategies

Having established that the cattle movement network facilitated widespread epidemic establishment, we used counterfactual simulations to evaluate the degree to which earlier or stronger interventions might have mitigated the outbreak (Fig. S22). The calibrated simulation model with the federal order at its actual implementation date resulted in a median of 510 infected counties (50% confidence interval [CI]: 330–716; 95% CI: 106–1,230), out of 2,504 counties with at least one dairy premises in the network realizations used for our epidemic model. Without any interstate intervention, this rose to 755 counties (50% CI: 458–1,059; 95% CI: 136–1,488; Fig. 5a,b).

**Figure 5.**
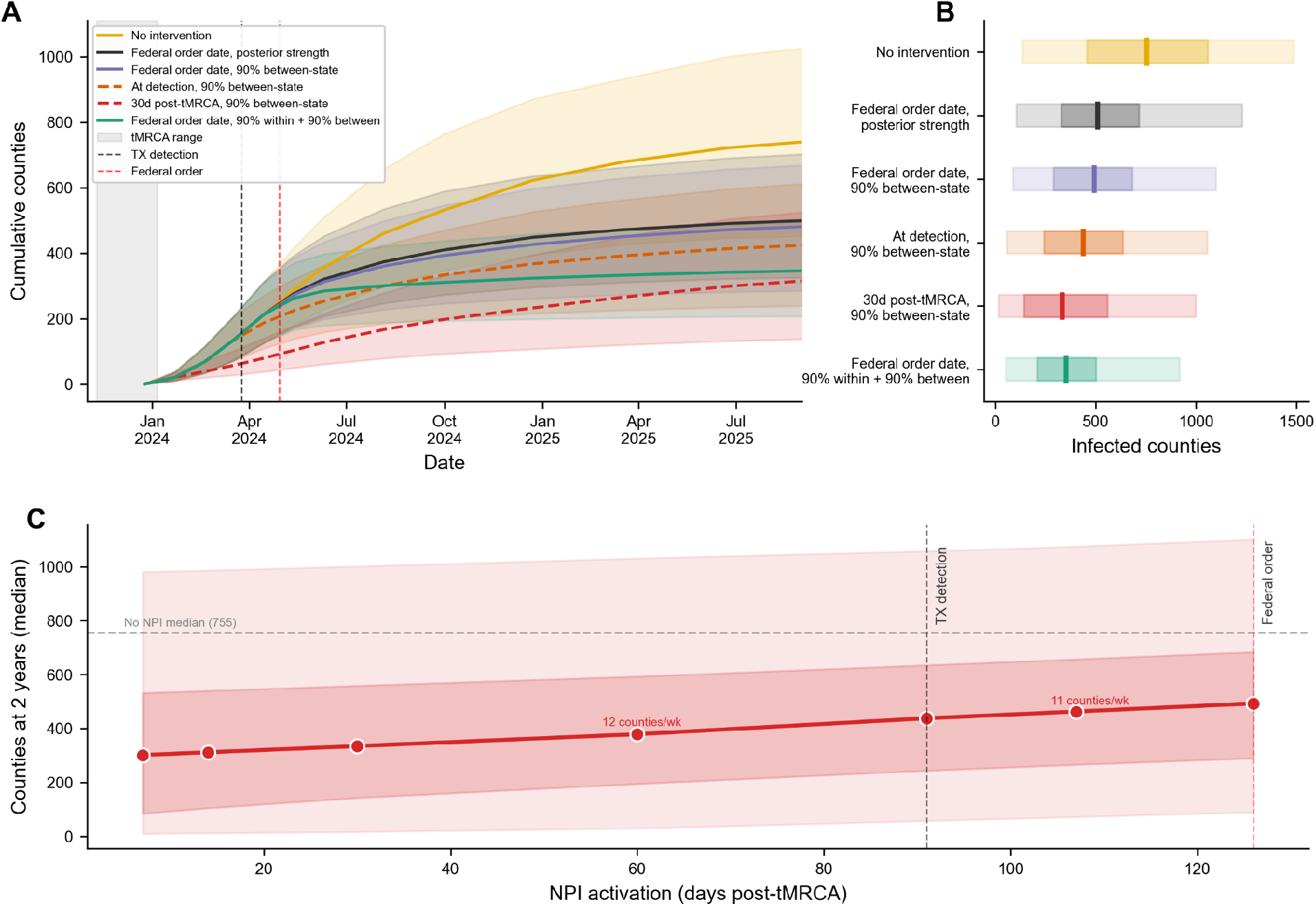
Effects of different intervention strategies on the trajectory of the outbreak. **a**, Cumulative counties infected in the simulations under different intervention strategies: no NPIs (yellow), the empirical between-state intervention (black), a between-state intervention with 90% effectiveness deployed on the same day as the actual federal order (purple), a between-state intervention with 90% effectiveness deployed on day of first detected case (orange), a between-state intervention with 90% effectiveness deployed 30 days after the tMRCA of B3.13 in dairy cattle (red), and a within- and between-state intervention with 90% effectiveness deployed on the same day as the actual federal order (green). tMRCA, date of earliest detection, and date of federal order indicated by a grey bar, a dashed black line, and a dashed red line, respectively. Median and interquartile ranges are indicated. **b**, Number of counties infected at the end of the simulations (1 September 2025) across the six intervention strategies described in (a). Dark line, median; dark rectangle, interquartile range; light rectangle, 95% confidence interval. **c**, Number of counties infected at the end of the 2-year simulation plotted against when the standardized between-state non-pharmaceutical intervention was implemented, with the increased average number of infected counties per week of delay indicated. Dots indicate dates at which non-pharmaceutical interventions were implemented. Figure S23 shows the numbers of counties and states infected for all counterfactual scenarios.

To examine whether a stronger interstate policy could have reduced spread, we standardized the policy effect such that interstate viral transmission is reduced by 90% (corresponding to a non-pharmaceutical intervention (NPI) factor of 0.1; the full distribution of inferred NPI values is shown in Fig. S15), which resulted in a slight decrease in the number of counties infected (median of 493 infected counties; 50% CI: 290–684; 95% CI: 90–1,101). Implementing the same intervention on the day of the earliest outbreak detection (25 March 2024), rather than at the actual federal order date, reduced the median to 438 infected counties (50% CI: 244–635; 95% CI: 58–1,057) (Fig. 5b). This modest reduction reflects that, by the time any interstate policy could have been enacted following detection, many critical states would have already been seeded (Fig. S23, Table S7).

A more effective interstate intervention would have required implementation before the outbreak was detected. Had surveillance identified an infected premises within 30 days of the tMRCA (Fig. S22)—approximately two months before the first outbreak was confirmed—and a policy implemented immediately that resulted in a 90% reduction in interstate transmission, the epidemic would have been limited to a median of 335 counties (50% CI: 143–559; 95% CI: 17–1,001), a reduction of ∼32% relative to the standardized interstate policy (Fig. 5b).

However, the federal order targeted only interstate movement. Because within-state cattle movement drove viral persistence (Fig. 3) and likely sustained the epidemic through much of 2024 and 2025 (Fig. 2a), we examined whether complementary within-state restrictions could have further reduced spread. Applying an intervention that resulted in a 90% reduction of within-state, between-county viral transmission on top of the standardized interstate policy, and beginning on the same day as the actual federal order, reduced the median number of infected counties from a median of 493 to 353 (50% CI: 209–502; 95% CI: 53–918). This change resulted in a 28% reduction in infected counties, here attributable to limiting within-state amplification.

More broadly, the relationship between implementation timing and epidemic size was approximately linear: each week of delay in policy implementation was associated with an additional median 11–12 infected counties per week (Fig. 5c). Together, these counterfactual analyses suggest that the trajectory of the B3.13 epidemic was shaped less by the strength of the policy response than by the interval between viral introduction and its detection.

Lastly, because the Texas Panhandle was the likely epicentre of the epidemic, we examined whether a rapid, intensive, but more precisely targeted containment strategy could achieve greater epidemic control while requiring intervention in fewer locations than the broader federal movement order. To isolate the effects of targeting rather than intervention strength, we compared idealized interventions that completely prevented between-county transmission and differed only in the counties targeted and when the interventions were implemented (Fig. S24, Fig. S25, Table S7).

Geographically-defined containment zones (ranging from the 14 dairy counties in the Texas Panhandle to progressively larger rings encompassing >400 counties) only achieved modest reductions, even when implemented just 14 days after the start of the outbreak (Fig. S25). By contrast, targeting counties connected through the cattle movement network was substantially more effective. Restricting movement from the 41 counties reachable via one shipment from the Panhandle resulted in similar reductions to targeting all of Texas and six other states (Fig. S24). Targeting the 102 counties within two links of the Panhandle in the cattle movement network reduced the epidemic to a median of 12 affected counties, compared with the 430 affected counties for the similarly sized geographic buffer. Although delaying implementation to 30 days after the start of the outbreak diminished efficacy, the two-link network perimeter still reduced the epidemic to a median of 382 affected counties, ∼25% fewer than under the federal order. These results demonstrate that network-informed targeting can substantially outperform geographically-defined containment, particularly when interventions are implemented early in an outbreak.

## Discussion

We find that the spread of H5N1 B3.13 in dairy cattle, from its probable emergence in the Texas Panhandle to 16 states across the US, was driven primarily by cattle movement. Although the federal order restricting interstate cattle movement substantially reduced interstate viral lineage movement, extensive within-state spread persisted, with internal cattle movement and network density predicting prolonged transmission following introduction. Our simulations across the dairy cattle movement network further indicate that the connectivity of the outbreak epicentre facilitated the widespread dissemination of B3.13 and that earlier, network-informed interventions would have been considerably more effective than broader interventions implemented after widespread cryptic transmission.

Wild birds, particularly ducks and other waterfowl, are the predominant natural reservoir of influenza A virus genetic diversity^17^. Despite the emergence of B3.13 in dairy cattle being due to a spillover of B3.13 from wild birds, and despite its initial geographic expansion among dairy cattle coinciding with the main northward migratory period of waterfowl^18,19^, we found no evidence that wild birds contributed to the geographic spread of B3.13 among dairy cattle. Our results indicate there was recurrent transmission from dairy cattle into wild birds—predominantly peridomestic species (*e.g.*, robins, grackles) that share habitat with farms—but little evidence of onward transmission among them. Given the devastating effects of the H5N1 panzootic on wild bird populations globally^20,21^, the impact of cattle-origin viruses on wild birds has been limited, suggesting that many wild bird populations may already have substantial population immunity, that B3.13 is not a particularly fit genotype in wild birds, or that the peridomestic species infected with B3.13 may be less efficient vectors for transmission to other birds.

The movement-driven spread of B3.13 reflects broader structural changes in the US dairy industry over recent decades that have shifted production toward fewer, larger operations, with geographically dispersed sites connected by extensive cattle movements^22–25^. Our findings provide empirical evidence that this increasingly interconnected network structure shaped epidemic dynamics across multiple spatial scales. Cattle movement consistently explained interstate viral spread more strongly than geographic proximity, while county-level network connectivity predicted both multi-state spread and prolonged transmission within states. Although the sustained transmission of H5N1 in dairy cattle was unprecedented, the structural conditions that facilitated its dissemination may be becoming more common. Intensification and consolidation of dairy production are occurring across many regions of the world, including South America, South and Southeast Asia, and parts of Africa^26^. As livestock production systems become more interconnected, pathogens that successfully establish within these networks may be more likely to cause large-scale epidemics. Concomitantly, shifting network structure, and variable movements through it, could also lead to less predictable and sub-exponential growth, as well as variable geographic dynamics, emphasizing the need to document how the network changes over time^27,28^.

The predominance of cattle movement over geographic proximity in explaining B3.13 spread is consistent with a broader pattern of infectious disease transmission. Previous theoretical work has demonstrated that specific network structures can determine whether newly emerged pathogens establish sustained transmission^29,30^. Livestock movement networks have been implicated in the spread of influenza among poultry premises in France^31,32^ and China^33^ and swine farms in the US and globally^34–36^, and human mobility networks have been shown to explain pathogen dispersal more accurately than geographic proximity for both SARS-CoV-2^37,38^ and seasonal influenza^39^. Our findings extend these observations to another large-scale livestock epidemic, demonstrating that movement networks shaped epidemic dynamics across multiple stages of epidemic spread, from county-level establishment to prolonged transmission and interstate dissemination.

Our findings also echo the 2001 foot-and-mouth disease epidemic in the United Kingdom, where nationwide movement bans reduced the reproduction number but were insufficient to interrupt transmission without additional ring culling around infected premises^40,41^. Although the US federal order requiring pre-movement testing for interstate dairy cattle transport reduced the interstate spread of B3.13, extensive within-state transmission persisted. Notably, the largest outbreak occurred in California, where B3.13 was introduced only after implementation of the federal order, illustrating how even relatively few introductions could give rise to substantial local outbreaks. In both epizootics, long-range livestock movements seeded new outbreaks, whereas local transmission sustained them, illustrating that interventions targeting only a single spatial scale are unlikely to be sufficient. Further, interstate transmission appears to have persisted into 2026 despite the testing policy. Determining how interstate spread has persisted is a priority, as ongoing movement may be sustaining an epidemic that otherwise might have gone extinct.

The movement network model used here is both predictive and explanatory with respect to B3.13 transmission across the United States, despite being parameterized using historical cattle movement data collected years before the emergence of H5N1 in dairy cattle^11^. Although contemporary, high-resolution movement data would undoubtedly improve inference and should remain a priority for surveillance, our results demonstrate that historical movement patterns nevertheless captured sufficient structural information to explain and predict epidemic dynamics. The resulting county-level risk map identified locations predisposed to seed and sustain large outbreaks, suggesting that network-informed risk maps could prioritize surveillance for future pathogens entering the dairy cattle movement network. Our findings reinforce the importance of publicly available livestock movement data for epidemic preparedness^41^ and suggest that epidemic preparedness should integrate pathogen-agnostic surveillance with an understanding of the structure and dynamics of movement networks.

The B3.13 epizootic has continued to evolve beyond the emergence and early dissemination period examined here. There appears to have been a recent shift in the geographic distribution of outbreaks, with fewer occurring in California and more in Idaho and Utah, as well as a resurgence in Texas^3^. However, the limited availability and temporal and geographic sampling biases in viral genomic data (*e.g.*, no genomes from Utah or Texas in 2026 despite outbreaks occurring there) and lack of associated metadata make it challenging to determine whether the epidemiological processes identified here continue to shape viral spread as the epidemic progresses. These surveillance limitations also prevent us from resolving the mechanistic routes of H5N1 transmission among dairy cattle, the within-premise and local between-premise dynamics that sustain spread, and whether these processes have changed as the epidemic has evolved.

Between-state policies would likely have been more effective had they been implemented earlier, but this would only have been possible with more timely surveillance and rapid granular reporting of detections. Our counterfactual analyses indicate that the interval between viral emergence and detection was more impactful in limiting viral spread than the strength of the intervention alone. The rapid identification of the H5N1 2.3.4.4b D1.1 genotype through bulk milk testing^42,43^ illustrates how enhanced surveillance can detect emerging pathogens before they become widespread, even when it remains uncertain whether the detected pathogen will establish sustained transmission. Further, given the substantial host range of lineage 2.3.4.4b—now spanning birds and mammals^44^—influenza A virus should be considered a candidate pathogen during outbreaks of unknown aetiology, including in species not previously recognized as susceptible. Real-time effluent or wastewater monitoring could reduce detection times further still, potentially to days rather than weeks or months. In the absence of widespread vaccination, strengthening surveillance systems capable of detecting both successful and unsuccessful pathogen emergence events will be essential for limiting future epidemics. Coupled with an understanding of livestock movement networks, earlier multi-modal^45^ and pathogen-agnostic surveillance offers the greatest opportunity to prevent the establishment and large-scale dissemination of emerging pathogens before they become a substantial risk to animal and human health.

## Methods

### Dataset curation

An H5Nx clade 2.3.4.4b dataset for North America, South America and Antarctica was compiled by querying the Global Initiative on Sharing All Influenza Data (GISAID)^46^ and NCBI Virus^47^ databases for all H5Nx clade 2.3.4.4b samples collected between 1 November 2021 – 9 January 2026 with available genomic sequences for all eight gene segments (accessed on 9 January 2026). Additionally, consensus genomes from raw reads deposited to the NCBI Sequence Read Archive and associated with BioProjects PRJNA1102327, PRJNA1122849, PRJNA1134696, PRJNA1219588, PRJNA1207547, PRJNA980729 were downloaded from the Andersen Lab’s avian-influenza github repository (https://github.com/andersen-lab/avian-influenza). The samples in each dataset were assigned a genotype using the GenoFLU genotyping tool^48^. A sequence dataset for genotype B3.13 (n = 4,516) was compiled for each gene segment (i.e., PB2, PB1, PA, HA, NP, NA, MP and NS) after removing duplicate sample entries across databases. We subsequently excluded all virus genomes sampled before March 2024 (when the outbreak started), B3.13 genomes sibling to the dairy cattle outbreak clade, genomes sampled from an individual emu farm, genomes associated with a pet food outbreak in Oregon, and genomes with incomplete dates or unknown locations. Each segment dataset was aligned using MAFFT v.7.526 and trimmed to the coding sequence region^49^. All eight segment alignments were concatenated into a single sequence per sample using ConcatenatorGUI v0.3.1 (ref.^50^). Accessions for all genomes used are available in Table S8.

Outbreak data were downloaded from https://www.aphis.usda.gov/livestock-poultry-disease/avian/avian-influenza/hpai-detections/hpai-confirmed-cases-livestock (last accessed 17 February 2026). We retained only confirmed B3.13 cattle detections, excluding poultry outbreaks and D1.1 genotype cases. World Animal Health Information System (WAHIS) outbreak records were matched to USDA detections using USDA event identifiers and state-level information. WAHIS, maintained by the World Organisation for Animal Health (or WOAH), is a global platform for reporting official animal disease information, including outbreaks and other epidemiologically significant events in domestic and wild species. WAHIS outbreak reports (accessed through https://wahis.woah.org/#/event-management) provide event-level information submitted by national veterinary authorities, including outbreak location, disease, report date, event status, and relevant epidemiologic dates.

Dairy industry data used for the phylogeographic analyses (see below) were obtained from USDA NASS^10^ for 2024, including dairy cattle inventory, milk production per cow, number of herds, and number of dairy cows and calved cows per state. We calculated a replacement ratio — the proportion of calved dairy cattle relative to total dairy cattle — as a proxy for the rate of new animals entering the herd. Pairwise geographic distances between state centroids were calculated as great-circle distances.

One set of cattle mobility data was obtained from USAMMv3 (ref.^11^), comprising 1,000 stochastic network realizations of daily dairy cattle shipments among premises across the contiguous United States over one year. The model was fitted to interstate certificates of veterinary inspection (ICVI) data, which document livestock movements across state lines, from 2009 and county-level demography from the 2012 NASS census in a Bayesian framework, with shipment rates and sizes varying by premises type, herd size, and season.

A second set of state-level dairy cattle mobility data were obtained from Cabezas *et al.* (2021; ref.^51^). These data are derived from ICVI records from 1 April 2015 through 31 March 2016, collected from State Animal Health Officials in the 48 contiguous US states; Alaska and Hawaii were not included (Table S2). An initial dataset comprised 735 directed state-to-state links among 36 states, representing 60,466 shipments in total. An additional 411 links representing 8,641 shipments were inferred for unobserved states missing from the initial dataset based on the 2012 Census of Agriculture and the reported values from counties and states surrounding the missing states. Each directed link represents movement from an origin state to a destination state weighted by the number of shipments. We used the full dataset, comprising 1,146 directed links among all 48 contiguous states (69,107 shipments). These two mobility datasets represented the most recent nationwide cattle mobility datasets available at the time of analysis.

### Phylodynamic and phylogeographic inference of B3.13 in dairy cattle, domestic avians, and wild avians

We first subsampled our larger dataset to make our analysis computationally tractable by randomly selecting up to 25 genomes for every month-state-host combination from March 2024 onward, assuming dairy cattle, wild bird, and captive bird hosts. We then jointly inferred the genealogy and the host state of each lineage of the resulting 1,358 genomes under an asymmetric forward-in-time (FIT) discrete phylogeographic model^8^ with Bayesian Stochastic Search Variable Selection (BSSVS) implemented in BEAST X v10.5.0 (ref.^52^) with BEAGLE v4.1 (ref.^53^). We modelled sequence evolution using an HKY substitution model with 4 discretized-gamma categories^54^ and an uncorrelated relaxed clock with an underlying lognormal distribution^55^. We ran a chain of 100 million states, assessed convergence and mixing in Tracer v1.7.2 and discarded 10% of the chain as burn-in. We used the highest independent posterior subtree reconstruction (HIPSTR)^56^ approach in TreeAnnotator X to construct a time-calibrated summary phylogeny and catalogued host transitions across the post-burn-in posterior trees.

To evaluate whether the relative transition rates between hosts and the number of Markov jumps^57^ were robust to cattle sampling intensity, we repeated the FIT phylogeography while downsampling dairy cattle virus genomes. We randomly subsampled to 50%, 25%, and 12.5% of the original count, with each level a subset of the previous. All non-cattle virus genomes were retained, except at the 12.5% level where domestic avian virus genomes were also halved to maintain cattle as the majority host. We subsequently performed FIT phylogeographic inference for each subsampled dataset, using the same parameterizations as described above for the complete dataset.

We then applied the same model to reconstruct the geographic history underlying this host reconstruction, now defining a deme for each of the 16 states from which genomes were sampled and correspondingly estimating a 16-state asymmetric matrix of forward-in-time migration rates alongside a shared effective population size. Because the genealogy had already been informed by the host reconstruction, we did not re-estimate it; we instead fit the migration process to an empirical set of trees drawn from the posterior of the analysis above, resampling a tree during the chain so that phylogenetic uncertainty was propagated into the geographic inference. We ran a chain of 5 million states, assessed convergence and mixing in Tracer v1.7.2 and discarded 10% of the chain as burn-in.

### Phylogeography of B3.13 in cattle across 2024 and 2025 with state-level data

We first performed Bayesian phylogenetic inference using BEAST X v10.5.0 (ref.^52^) with BEAGLE v4.1 (ref.^53^) of 3,950 cattle and human virus sequences, with the human viruses considered to be sentinels for cattle surveillance and not markers of onward transmission. Because of the relatively low degree of resolution offered by the sequence data, our full probabilistic model specification focuses on simple model specifications. We modelled sequence evolution using a general time-reversible (GTR) substitution model^58^ with 4 discretized-gamma categories^54^, a strict molecular clock model with an informative normal prior on the evolutionary rate based on our previous analyses, and a normal prior on the root height based on our previous analyses^4^. As a coalescent tree prior, we used a non-parametric skygrid prior^59^ with 25 grid points and a cutoff of two years. We ran an MCMC analysis of 100 million generations, subsampling every 10 thousand iterations to continuous parameter log and tree files. Convergence and mixing were assessed in Tracer v1.7.2 (ref.^60^), and 10% of the chain was discarded as burn-in. All relevant effective sample size (ESS) values were >200. We subsampled the resulting posterior tree distribution to create an empirical tree set comprising 225 trees.

We used a discrete phylogeographic approach with a log-linear parameterization of the transition rates between locations as a function of potential predictors^9^, and tested the association of viral lineage dispersal with geographic, dairy cattle mobility, and dairy industry-related factors (see Table S1 for a complete list of tested predictors). We also tested the effect of sampling bias by including a predictor based on the residuals of a regression of the number of genomes against dairy inventory, outbreak counts, and number of herds in each state. The predictors were log-transformed and standardized after adding a pseudo-count to each entry when appropriate.

We first performed these analyses using time-homogeneous and time-inhomogeneous parameterizations including the predictors in Table S1 and with BSSVS. All of the predictors were time-invariable, except for the dairy cattle mobility data obtained from USAMMv3, a stochastic model of daily cattle movements fitted to Certificates of Veterinary Inspection (CVI) records from 2009 and comprising 1,000 network realizations. For the time-homogeneous parameterization, we used the sum of all movements between states across the whole year, averaged across all 1,000 network realizations, as the predictor. For the time-inhomogeneous parameterization, we used a previously developed epoch extension^61^ to specify weekly intervals and incorporated weekly mobility matrices based on the USAMMv3 data. In this parameterization, we used weekly sums of the movements between states, averaged across the 1,000 network realizations. The rest of the predictors were kept constant through each epoch. We assumed time-homogeneous effect sizes and inclusion probabilities, with indicator variables that model predictor inclusion.

After identifying the subset of predictors supported by the initial analyses (Fig. S5), we subsequently evaluated whether their effects differed across three epidemiologically defined time periods (“eras”): before the confirmed detection of the first B3.13 outbreak in dairy cattle on 25 March 2024 (“pre-detection”), between this first detection and the implementation of a federal order on 29 April 2024 restricting cattle interstate movement (“pre-response”), and after the implementation of the federal order (“post-response”). We again used a time-inhomogeneous parameterization with weekly epochs, except now assuming inclusion probabilities of 1.0. Additionally, we included era-specific rate scalars on the overall rate of geographic transitions to account for temporal variation in the overall rate of lineage movement, ensuring that log-linear coefficients reflect predictor effects rather than changes in the baseline transition rate. As a sensitivity analysis and to determine during which eras individual predictors were supported, we applied the same time-inhomogeneous parameterization while assuming time-homogeneous effect sizes and inclusion probabilities per era.

To efficiently draw inferences under these time-homogeneous and time-inhomogeneous parameterizations, we fit the spatial diffusion process to the set of empirical trees inferred under the simpler specification described above. Model and inference specifications for the different analyses are available as BEAST XML input files on GitHub (https://github.com/pekarj/h5n1_b313_phylogeography). Convergence and mixing were assessed in Tracer v1.7.2, and 10% of the chain was discarded as burn-in. All relevant ESS values were >200.

Subsequent downstream analyses were done using the results from the time-inhomogeneous inference with covariates with inclusion probabilities of 1.0 and the three rate scalars. Cataloguing of between-state transitions was done using the Markov jump histories^57^. We used the HIPSTR approach in TreeAnnotator X to construct a time-calibrated summary phylogeny from the posterior tree distribution after removing burn-in.

### Phylogeography of early B3.13 genomes with county-level data

We performed a FIT discrete phylogeographic inference using a flexible asymmetric^8^ 18-state continuous-time Markov chain model, without BSSVS, on 149 B3.13 genomes sampled from avian, dairy cattle, human, and other mammalian hosts from the early outbreak that had county-level metadata available. We considered the non-cattle genomes to be sentinels for cattle spread and potentially indicative of where the outbreak could have originated. We modelled sequence evolution using a GTR model with 8 discrete-gamma categories, a strict molecular clock model, and an exponential growth coalescent tree prior. Because of the low sequence diversity so early in the outbreak, we used an informative normal distribution prior on the evolutionary rate based on our larger analysis of 3,950 genomes. We ran a chain of 100 million generations, subsampling every 10 thousand iterations to continuous parameter log and tree files. Convergence and mixing were assessed in Tracer, and 10% of the chain was discarded as burn-in. All relevant ESS values were >200. We used the HIPSTR approach in TreeAnnotator X to construct a time-calibrated summary phylogeny from the posterior tree distribution after removing burn-in.

All phylogenetic trees were processed using TreeSwift v1.1.42 (ref.^62^) and visualized using baltic v1.0.0 (ref.^63^)

### Within-state persistence

To quantify within-state viral persistence, we estimated the total phylogenetic branch time per state by averaging over the last 225 posterior trees from the time-inhomogeneous state-level phylogeographic analysis with three predictors and era-specific rate scalars. For each posterior tree, every branch was allocated to one or more states based on the discrete state annotations and Markov jump history^57^. Branches with no recorded state transitions were assigned entirely to the child node’s annotated state. Branches with one or more transitions were split at the sampled transition times, with each segment allocated to the corresponding state. Total branch time per state was computed as the sum of all allocated branch segments across the entire tree. The number of introductions per state was also estimated from the Markov jump history, with each recorded transition into a state counted as one introduction event. Texas was assigned an additional introduction to account for the root origin. Branch time per introduction was then estimated as the ratio of total branch time to the number of introductions for each state.

For each state included in the phylogeographic analysis, we computed network summaries from the 1,000 USAMMv3 cattle movement network realizations. These realizations represent movements between individual premises. Because the network data predate the epidemic by approximately 15 years (2009 versus 2024–2025), individual premises may have entered or exited the network over time. To reduce sensitivity to this premise-level turnover, we first aggregated shipments to the county level before calculating network summaries. Daily county-to-county shipments were aggregated annually, then summarized per state. Internal network density was computed as the number of realized within-state shipping routes divided by the maximum possible directed edges between dairy counties in that state. Internal shipping volume was the total number of cattle moved in within-state routes. Incoming routes is the number of interstate shipping routes directed into each state. Incoming shipment count is the number of cattle shipped into each state. All features were averaged across the 1,000 network realizations. The number of dairy counties per state, herd density (dairy cattle per km²), and annual replacement ratio were derived from 2024 USDA NASS data.

We examined association between these seven predictors and state-specific total branch time and branch time per introduction in a post hoc fashion. We first assessed collinearity among the predictors using Spearman rank correlations on their log-transformed values. Given that several predictors were strongly correlated and the limited number of states (n=16), we fit univariate ordinary least squares regressions for each predictor on each posterior tree sample independently, with both predictor and response log-transformed and the log-predictor standardized prior to regression. For each posterior tree sample, we tested whether the slope coefficient differed from zero using a t-test, and reported the distribution of R² and β coefficients across posterior samples. Robustness was evaluated by a state-based leave-one-out analysis. As there were substantially more genomes from California than any other state, we performed a separate sensitivity analysis subsampling taxa from California at 50%, 25% and 10%, pruning the posterior tree samples accordingly, and recomputing branch times and introduction counts from the pruned trees before repeating the regressions.

### Epidemic model using dairy cattle networks

We developed a stochastic susceptible-infected-recovered (SIR) network model to simulate H5N1 B3.13 spread among US dairy cattle at the county level. Each US county with at least one dairy premise in the USAMMv3 network realizations (n = 2,504 counties) was represented as a single node with a binary infection state. Inter-county cattle shipments were drawn from USAMMv3, which comprises 1,000 independent network realizations, each providing a full year of daily directed edge lists specifying the number of cattle shipped between county pairs.

Early epidemiological reports suggested that the epidemic could have begun in Castro County, Texas, and the phylogeographic reconstruction of B3.13 using the early genomes with county-level metadata inferred the location of the most recent common ancestor to be Castro County (posterior probability=0.68). Our simulations were therefore seeded in Castro County. Simulation day 0 corresponded to the tMRCA of B3.13 in cattle, estimated at 79–137 days before first detection in Texas (based on the 95% HPD of the tMRCA; ref.^4^). Each simulation day mapped to the corresponding day of the year in the movement network.

On each day in the simulation, for each infected county *u* with a shipment edge to susceptible county *v*, transmission occurred with probability

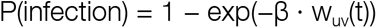

where β is a per-head transmission coefficient and w_uv_(t) is the number of cattle shipped from *u* to *v* on day t. That is, larger cattle shipments have a higher probability of transmitting infection, with the probability approaching 1 as shipment size increases. Each infected county recovered after a fixed duration scaled by local herd characteristics:

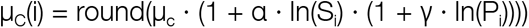

where μ_c_ is a base recovery period (days), S_i_ is the number of dairy cows in county i, P_i_ is the number of dairy premises, and α and γ are scaling parameters. No reinfection was permitted.

The federal order of 29 April 2024, which required pre-movement testing for interstate cattle shipments, was modeled by reducing the volume on interstate edges by a factor f_NPI_ on simulation days at or after the order date, while leaving intrastate edges unaffected:

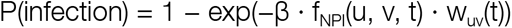

This design reflects the scope of the federal order, which regulated interstate but not intrastate cattle movement.

Each simulation was run for 830 days from the inferred tMRCA, yielding endpoints between February 2026 and April 2026 depending on the sampled tMRCA offset.

### Epidemic model calibration

We calibrated the six SIR parameters using approximate Bayesian computation with sequential Monte Carlo (ABC-SMC)^64^. Uniform priors were placed on all parameters: the per-head transmission coefficient β ∈ [0.001, 0.05], herd size scaling α ∈ [0, 1], premises scaling γ ∈ [0, 1], base recovery period μ_c_ ∈ [7, 59] days, interstate NPI factor f_NPI_ ∈ [0, 1], and tMRCA offset Δ_tMRCA_ ∈ [79, 137] days, where the tMRCA offset bounds correspond to the 95% HPD from molecular clock analysis. The model was calibrated against state-level outbreak data for 16 states with confirmed B3.13 outbreaks, using first detection timing per state and the rank ordering of county-level outbreak counts (Table S3).

The ABC-SMC distance function combined five components: (1) 1 minus the Spearman rank correlation between simulated and observed state arrival order, (2) log-scale RMSE of arrival timing for states detected more than 30 days after Texas, (3) a one-sided penalty for states with confirmed outbreaks within 30 days of the first confirmed Texas outbreak (25 March 2024) that were infected later in the simulation than observed, penalizing lateness more heavily for the earliest-detected states, (4) a fixed penalty for each observed state not infected in the simulation, and (5) 1 minus the Spearman rank correlation of county-level outbreak counts per state.

ABC-SMC proceeded over four rounds of 1,500 proposals each. In round 1, parameters were sampled from the prior; in subsequent rounds, accepted particles from the previous round were perturbed using Gaussian kernels with scales set to twice the standard deviation of accepted values (floored at 1% of the prior range), with reflection at prior boundaries. Each proposal was evaluated across 3 randomly selected network realizations with 2 simulations per network, and the top 5% of proposals were retained in each round, yielding 75 accepted parameter sets for downstream analyses.

### Epidemic simulations

For the primary simulations, we used Castro County, Texas, as the starting county for the epidemic. Using the 75 accepted parameter sets from ABC-SMC, we ran simulations across all 1,000 USAMMv3 network realizations with 5 independent simulations per network per parameter set, yielding 375,000 simulations per scenario. Simulations incorporated the observed federal order, with interstate movement restrictions activated 35 days after first detection in Texas and f_NPI_ drawn from the calibrated posterior. For each simulation, we recorded complete county-level transmission logs (source county, destination county, simulation day) and infection and recovery times per county, from which state-level arrival timing, outbreak size, and interstate versus intrastate transmission counts were computed.

To assess how the geographic origin of the epidemic influenced its trajectory, we seeded simulations from each of the 2,504 counties with dairy cattle across the 48 contiguous states in the USAMMv3 networks. For each county, the epidemic was seeded independently using all 75 ABC-SMC parameter sets across 200 network realizations with 5 simulations each, resulting in 75,000 simulations per county. The simulations were run with the federal order NPI implemented based on the ABC-SMC parameter sets. Multi-state spread probability was defined as the fraction of simulations in which the epidemic spread to at least 2 other states.

We then evaluated four sets of counterfactual scenarios. First, we performed a counterfactual with no interstate restrictions (f_NPI_ = 1). Second, we swept the strength of the federal order at the observed timing (day 35), fixing f_NPI_ at 0.1, 0.3, 0.5, 0.7, and 0.9. We subsequently used an f_NPI_ of 0.1 as a standardized policy for consistent comparison in subsequent simulations. Third, we varied the timing of a standardized federal order, implementing it at either 7, 14, 30, or 60 days post-tMRCA, on the day of confirmed detection of the first dairy cattle outbreak, or on 16 days after the confirmed detection of the first dairy cattle outbreak. The date of confirmed detection varies depending on the date of the tMRCA and is simulation-specific, but results in the equivalent of 25 March 2024 in the simulations. Lastly, we introduced within-state interventions that reduced intrastate cattle movement volumes by the same mechanism as the interstate restriction, applied independently to within-state edges. Within-state scenarios reduced intrastate transmission at the federal order date, testing a 90% intrastate reduction in addition to the standardized 90% interstate restriction (*i.e.*, the intervention restricts intercounty transmission across the country).

To evaluate whether targeted local containment could have mitigated the epidemic, we simulated restricting outbound cattle movement from progressively larger containment zones centered on the outbreak epicentre in the Texas Panhandle. Since fewer regions would be impacted than with the federal order, complete prevention could be more realistic. Six containment perimeters were tested: the Texas Panhandle (there are 26 counties in the Texas Panhandle, but only 14 have dairy cattle); the Panhandle plus two rings of geographically adjacent counties (47 dairy counties assuming Rook contiguity); the Panhandle plus four geographic rings (111 dairy counties assuming Rook contiguity); the Panhandle plus its top shipping destinations identified from USAMMv3 network data (41 dairy counties across 8 states, defined as counties receiving direct shipments from the Panhandle in at least 99% of network realizations); the Panhandle plus conditional two-hop network destinations (102 dairy counties across 14 states, where a second-hop county was included if the probability of receiving cattle from that county’s intermediary - given that the intermediary received cattle from the Panhandle - exceeded 99%); and all Texas dairy counties plus the dairy counties in 6 nearby states (NM, OK, AR, LA, KS, CO; 437 dairy counties). Each perimeter was tested at four policy implementation times: 14, 30, and 60 days after the tMRCA (the start date of the outbreak in the simulations), and at the date of first detection in Texas. All scenarios applied a 100% reduction in outbound movement volume from counties within the containment zone.

### County-level network features

For each seed county used in the alternative origin analysis, we computed network features from the USAMMv3 realizations, averaged across all 1,000 networks. We computed both static and temporal metrics.

Static metrics were computed from annual aggregate directed graphs: out-degree (number of unique destination counties), in-degree (unique source counties), out-volume and in-volume (total cattle shipped), PageRank, betweenness centrality, eigenvector centrality (computed on the undirected graph), closeness centrality, and clustering coefficient (undirected).

Temporal metrics captured the dynamic structure of shipment patterns over the year. Shipping consistency was defined as the fraction of days in the year on which a county shipped cattle. Maximum shipping gap was the longest interval (in days) between consecutive outbound shipments. Temporal PageRank was computed by averaging weekly PageRank scores across 52 weekly network snapshots. Temporal betweenness was the fraction of sampled time-respecting shortest paths passing through each county, where a path from A to B via C is valid only if the shipment from A to C occurs on or before the shipment from C to B. Temporal forward reachability was the number of unique counties reachable within 30 days via up to 3 hops along time-respecting paths.

These features were correlated with multi-state spread probability (1 minus die-out rate) across all seed counties to identify which network properties of the origin county most strongly predict whether an epidemic becomes established.

## Tables

Table S1. Covariates used for the phylogeographic analysis.

Table S2. Number of dairy cattle shipments between states based on ICVI records from 2015–2016.

Table S3. Earliest documented and confirmed infections of dairy cattle based on WAHIS and USDA, respectively.

Table S4. Predictors tested for association with the phylogenetic persistence time.

Table S5. Posterior calibration results for the epidemic simulation parameters.

Table S6. Static and temporal traits of individual counties based on the USAMMv3 network realizations.

Table S7. Epidemic outcomes after two years of simulated transmission across all counterfactual scenarios.

Table S8. Influenza genome accessions.

## Data and code availability

All genome accessions are available in Table S8. The XML files, code, and Newick trees are available at https://github.com/pekarj/h5n1_b313_phylogeography.

## Acknowledgements

JEP, MUGK and AR acknowledge funding from The Rockefeller Foundation (PC-2022-POP-005). JEP and AR acknowledge the support of the Wellcome Trust through grant 313694/Z/24/Z and MAS, AR and PL through grant 206298/Z/17/Z – “ARTIC network”. YS, PL, AR and MAS are supported by US National Institutes of Health (NIH) grant R01 AI153044. JOW and MAS are supported by NIH grant R01 AI192139. AMD and EP received funding from NSF grants 2210979 and 2210980. GD has received funding from the Research Council of Lithuania, Lithuania under the EMBO Installation Grant programme under grant No 5305. KG, KGA and MAS are supported by NIH grants U19 AI135995 and 3U19AI135995-07S2. GB and PL acknowledge support from the Internal Funds KU Leuven (Grant No. C16/26/012). GB acknowledges support from the Research Foundation - Flanders (“Fonds voor Wetenschappelijk Onderzoek - Vlaanderen,” G098321N), the European Union Horizon 2023 RIA project LEAPS (grant agreement no. 101094685), and the DURABLE EU4Health project 02/2023-01/2027, which is co-funded by the European Union (call EU4H-2021-PJ4) under Grant Agreement No. 101102733. T.P. was supported by by the UK Medical Research Council/Department for Environment, Food and Rural Affairs (Defra, UK) FluTrailMap-One Health consortium [MR/Y03368X/1], and by the BBSRC via the Pirbright Institute’s Strategic Programme Grants (ISPGs) [BBS/E/PI/230002A; BBS/E/PI/230002B], and BBSRC National Bioscience Research Infrastructure: Low Containment Services and Science Platforms grant [BBS/E/PI/23NB0003]. This work is supported in part by the Centers of Excellence for Influenza Research and Response, National Institute of Allergy and Infectious Diseases, National Institutes of Health (NIH), Department of Health and Human Services, under contract 75N93021C00014. This research was supported in part by the Intramural Research Program of the National Institutes of Health (NIH). The contributions of the NIH author(s) are considered Works of the United States Government. The findings and conclusions presented in this paper are those of the author(s) and do not necessarily reflect the views of the NIH or the U.S. Department of Health and Human Services, or the United States government. We thank Advanced Micro Devices, Inc., for their donation of massively parallel computing hardware. We gratefully acknowledge the authors and researchers from the USDA, public health agencies, originating laboratories and the submitting laboratories, who generated and shared through NCBI and GISAID the viral genomic sequences and metadata on which this research is based (Table S8).

## Competing interests

MAS receives contracts and grants from the US Department of Veterans Affairs, Johnson & Johnson and Gilead Sciences outside the scope of this work.

## Supplementary Note 1

To examine how the structure of the cattle movement network shaped the spatial trajectory of the B3.13 epidemic, we constructed a county-level transmission model calibrated to the observed outbreak. We first performed a phylogeographic analysis of an early subset of B3.13 genomes that had county-level metadata and inferred Castro County, Texas as the location of the most recent common ancestor (MRCA; Fig. S14). Given this inference and that the Texas Panhandle was the epicenter of the outbreak, we used Castro county as the starting point for our simulations. We initiated the epizootics at a previously inferred tMRCA (95% HPD: 9 November 2023–6 January 2024; ref.^4^) and incorporated the federal order restricting interstate dairy cattle movement on 29 April 2024 (see Methods; Fig. S15; Table S5). The calibrated simulations recapitulated the observed epidemic, with states that experienced outbreaks consistently infected in simulations and typically among the first to receive viral introductions (Fig. S16). After calibration, the number of states and counties affected were generally consistent across a range of start dates (Fig. S18), with a later start date necessitating a slightly greater transmission rate (Fig. S17).

**Figure S1.**
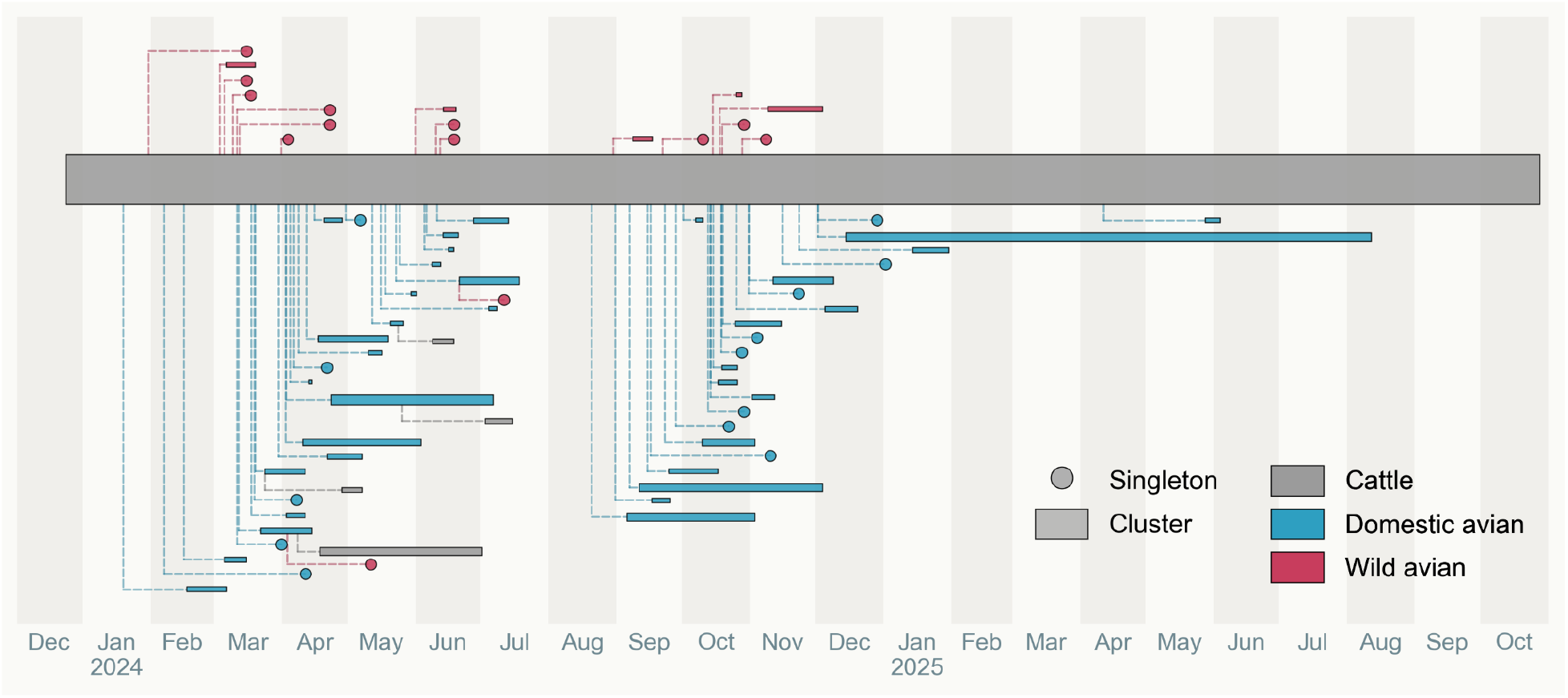
Frequent spillovers from dairy cattle to wild and domestic birds. Schematic of spillovers from the dairy cattle B3.13 outbreak, with the backbone and uninterrupted cattle transmission represented as the central grey rectangle, and each spillover from dairy cattle to other species visualized as circles (spillovers resulting in singletons) and rectangles (spillovers resulting in transmission clusters). Primary spillovers to wild avian and domestic avian species are positioned above and below the cattle backbone, respectively. Dashed L-shaped lines connect the cattle backbone to each spillover, originating at the time of the branch leading to the spillover. When the spillover leads to a single genome, the end of the branch is indicated by the location of the circle. When the spillover leads to a transmission cluster, the start (left side) of the rectangle is at the time of the most recent common ancestor of the transmission cluster, and the end (right side) of the rectangle is at the date of the most recent sample of the cluster. Subsequent spillovers from the transmission cluster follow the same pattern, except all said spillovers are located on the same side of the dairy cattle backbone as the initial spillover. However, these secondary spillovers are likely spillovers from dairy cattle, rather than birds, and appear to emanate from avian hosts due to unsampled dairy cattle virus diversity. Red represents wild avian hosts; blue represents domestic avian hosts; grey represents dairy cattle hosts. The heights of the rectangles representing the central dairy cattle outbreak and the spillover transmission clusters are proportional to their total branch lengths. Figure 1a shows the phylogeny with the spillovers indicated, and the full phylogeny is available via the online repository.

**Figure S2.**
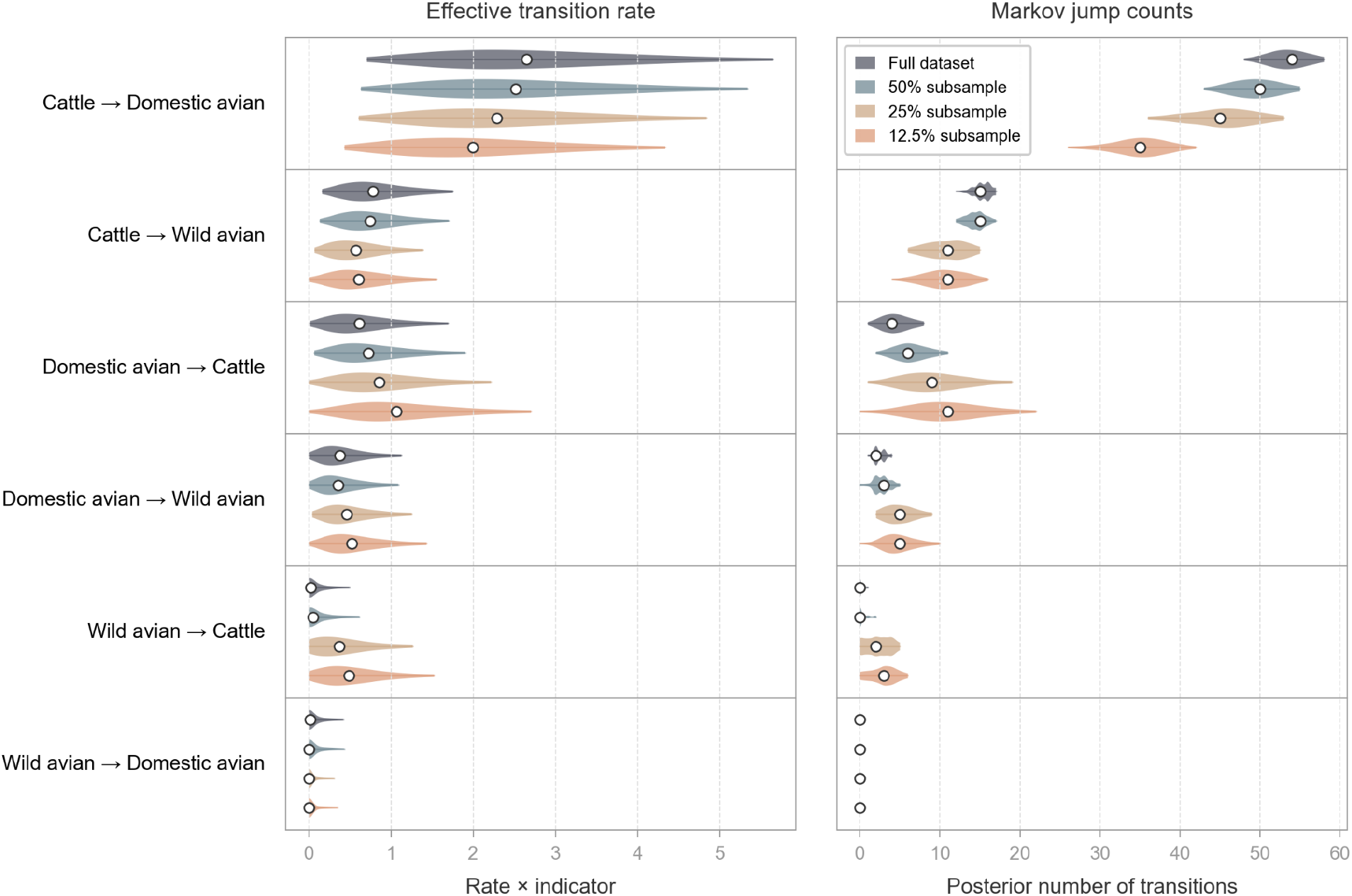
Host transitions occur predominantly from dairy cattle to other species. The effective host transition rate (left) and the posterior number of transitions between hosts (right), at different levels of subsampling of dairy cattle. The full dataset comprises 1,358 taxa (1102 dairy cattle viruses, 230 domestic avian viruses, 26 wild avian viruses), the 50% subsample comprises 807 taxa (551 dairy cattle, 230 domestic avian, 26 wild avian), the 25% subsample comprises 531 taxa (275 dairy cattle, 230 domestic avian, 26 wild avian), and the 12.5% subsample comprises 278 taxa (137 dairy cattle, 115 domestic avian, 26 wild avian; domestic avian were subsampled to 50% here to maintain dairy cattle as the majority host). Violins depict 95% highest posterior density (HPD), and white circles represent the median value.

**Figure S3.**
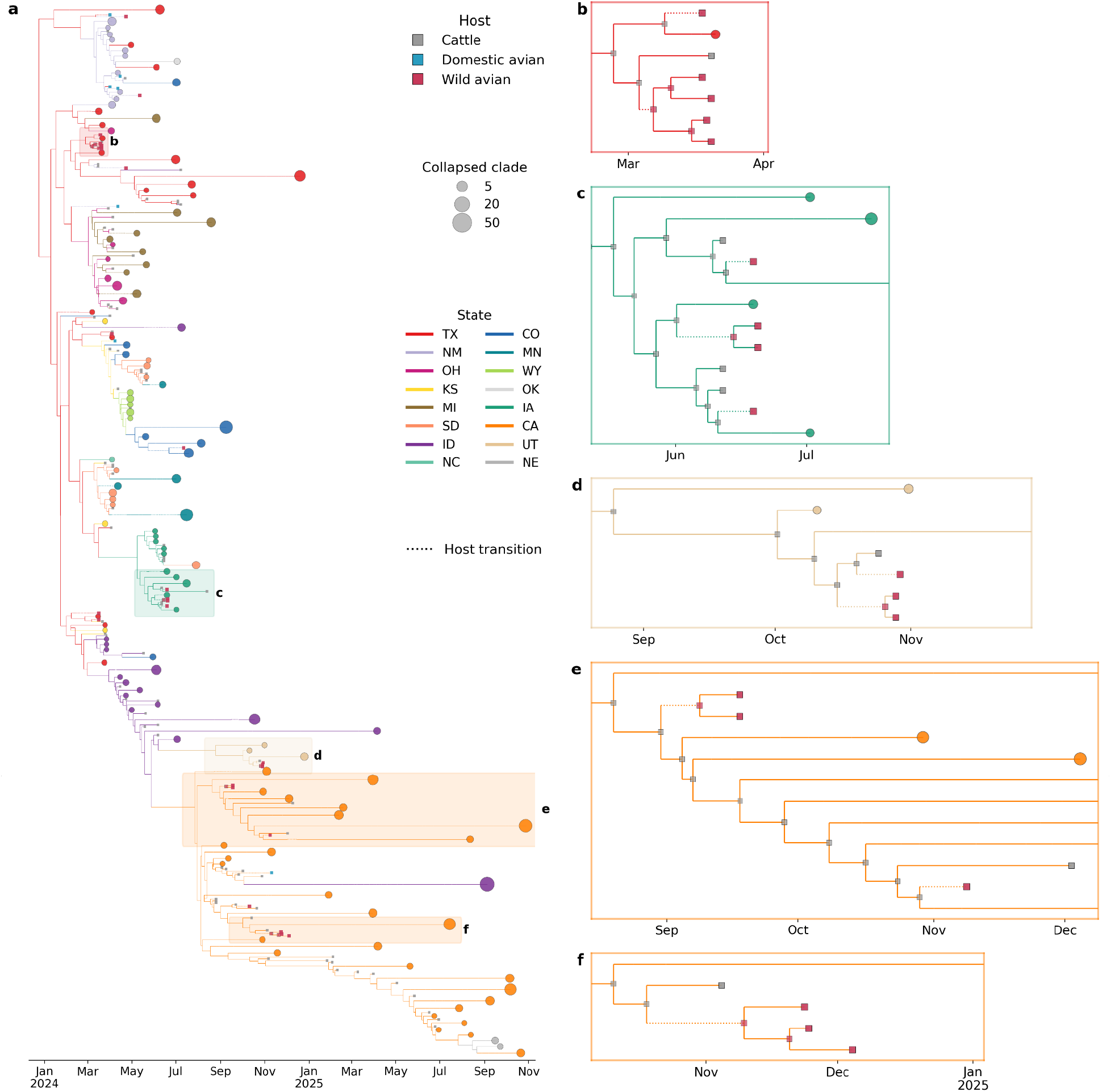
Wild avian transmission clusters represent within-state transmission. **a**, Time-scaled phylogeny of 1,358 H5N1 B3.13 sequences from a three-host phylogeographic analysis (cattle, domestic avian, wild avian). Monophyletic clades sharing a single location are collapsed into circles proportional to the number of constituent taxa, positioned at the date of the most recently sampled tip. Individual tips are shown where clades remain uncollapsed. All wild bird virus sequences are shown (i.e., no collapsed clade includes wild bird viruses). Branches and collapsed clades (circles) are colored by the most probable US state. Individual tips (squares) are colored by host. Dotted branches represent host transitions. Highlighted boxes indicate the subclades containing wild avian transmission clusters in Texas (**b**), Iowa (**c**), Utah (**d**), and California (**e**, **f**). Border color represents the state within which wild avian viruses circulated within the given subclade. All cattle-to-wild avian host switches and downstream wild avian transmission occur within the same geographic state. Trees within panels **b**–**f** truncated along the time axis for clarity.

**Figure S4.**
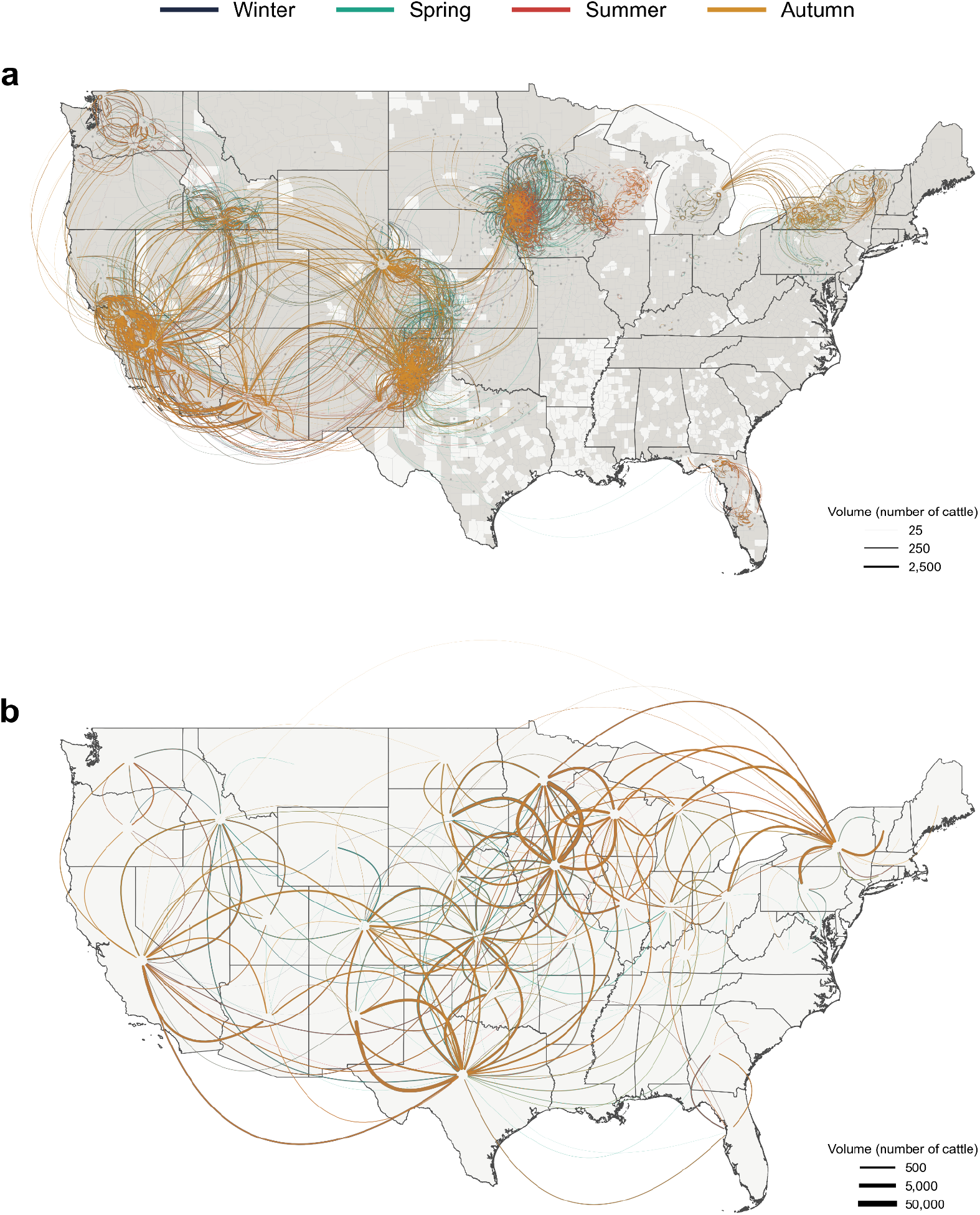
Dairy cattle mobility data. County-to-county (**a**) and state-to-state (**b**) dairy cattle shipments from the USAMMv3 network data, stratified by season and averaged across all 1,000 network realizations. Premise-level shipments were aggregated to counties (**a**) and states (**b**) before averaging. Arc thickness indicates average volume (head) shipped, and arcs curve clockwise from origin to destination. Shipments with fewer than 25 and 500 head of cattle are excluded from **a** and **b**, respectively, for clarity.

**Figure S5.**
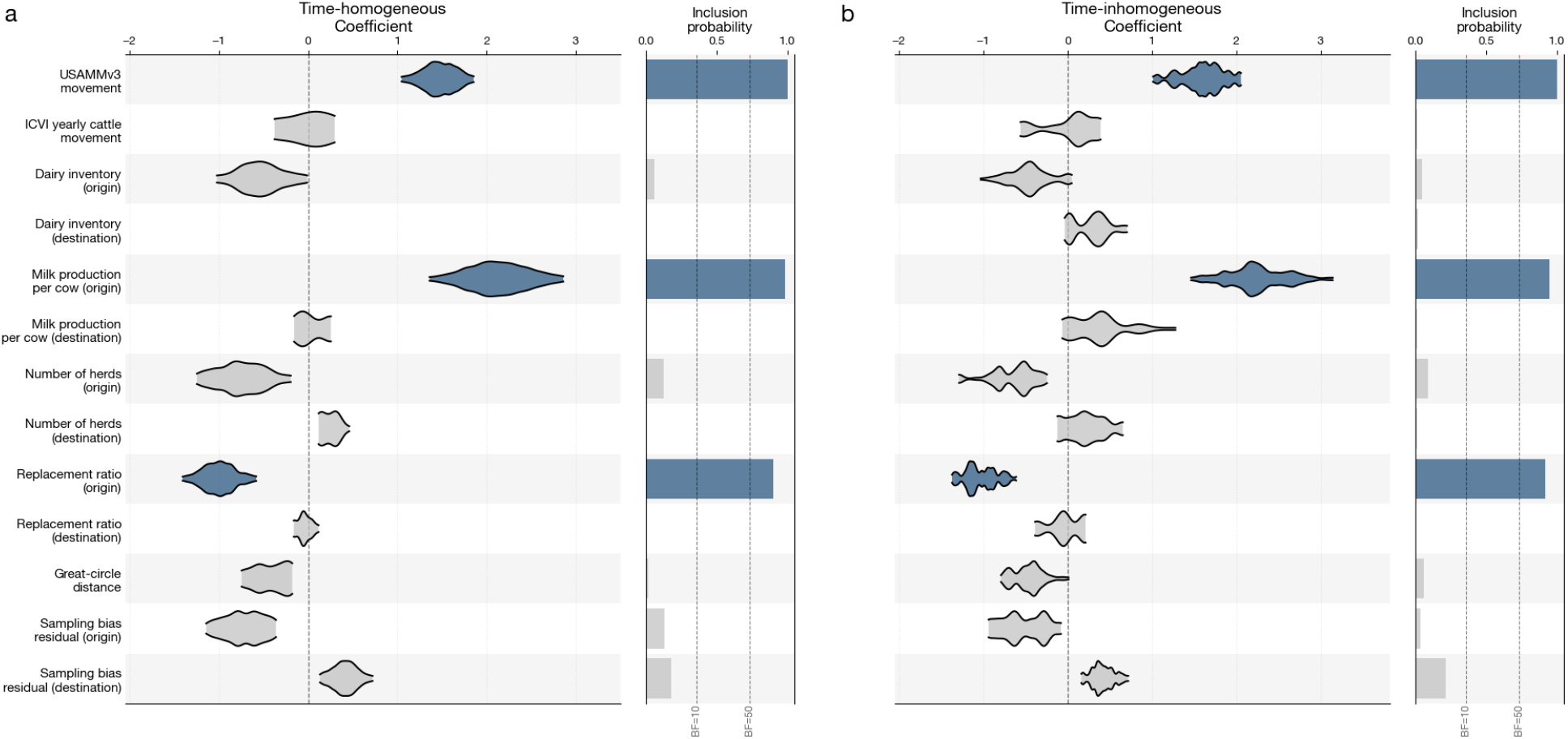
Predictors of H5N1 viral lineage movements in the US. Phylogeographic log-linear coefficients (left) and inclusion probabilities (right) for spatial spread covariates in (**a**) time-homogeneous and (**b**) time-inhomogeneous analyses. Dashed lines indicate Bayes factor thresholds of 10 and 50 in favor of covariate inclusion. Covariates with BF > 10 are highlighted in blue. Covariates are defined in Table S1.

**Figure S6.**
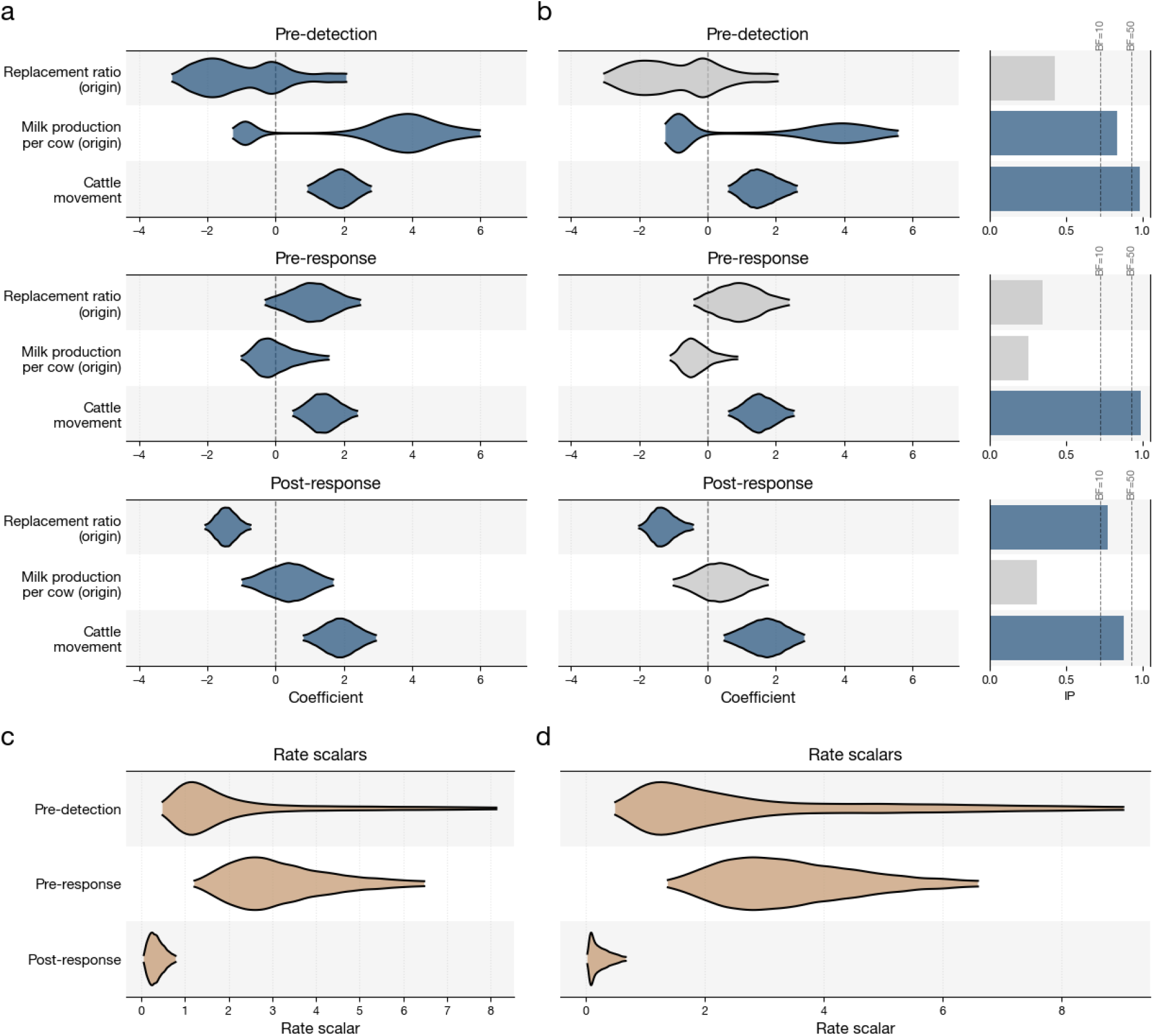
Supported predictors of H5N1 viral lineage movements across eras. **a–b,** Phylogeographic log-linear coefficients across the pre-detection (top), pre-response (middle), and post-response (bottom) eras for the three supported spatial spread covariates from the time-homogeneous analysis. **a**, Coefficients without inclusion probabilities. **b**, Coefficients with estimable inclusion probabilities, inferred via Bayesian stochastic search variable selection; corresponding inclusion probabilities and Bayes factor thresholds (dashed lines, BF = 10 and 50) are shown at right. **c–d**, Era-specific rate scalars for the models in **a** and **b**, respectively. Covariates without inclusion probabilities **a** and those with BF > 10 (**b**) are highlighted in dark blue. Covariates are defined in Table S1.

**Figure S7.**
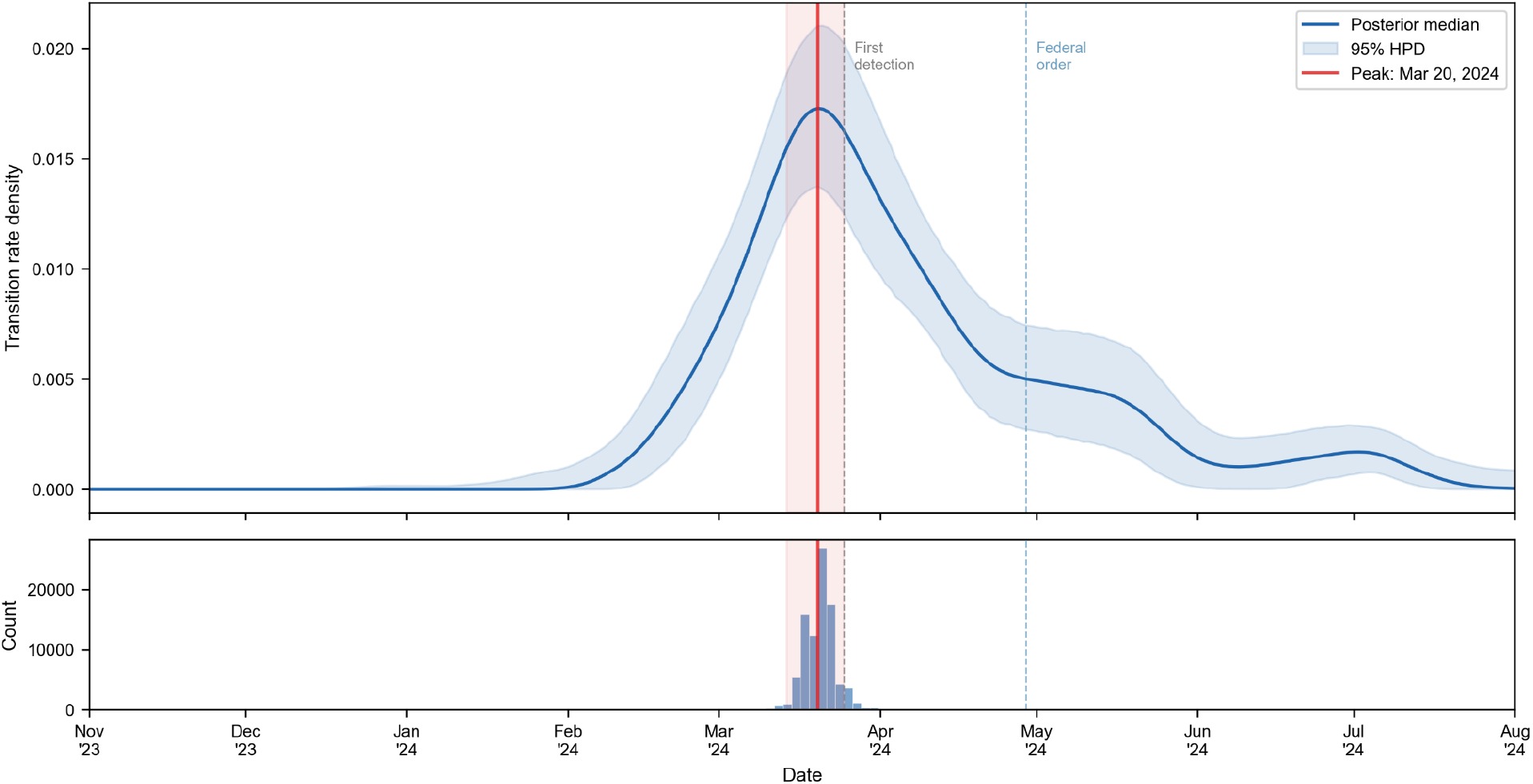
Peak timing of interstate viral transmissions. Upper panel: Gaussian kernel density estimate (σ = 7 days) of the interstate transition rate over time. The blue line shows the posterior median kernel density estimation (KDE) curve; the shaded band shows the 95% HPD interval across posterior samples at each time point. The red vertical line marks the posterior median peak date (20 March 2024), with shading indicating the 95% HPD interval (14 March—25 March). The grey dashed line marks first detection in Texas (25 March 2024); the blue dashed line marks the federal movement order (29 April 2024). The peak of interstate transmission precedes first detection by approximately 5 days, consistent with extensive cryptic spread driving the epidemic before surveillance detected it. Lower panel: Histogram of estimated peak times, showing the distribution of estimated peak timing. Each posterior sample’s jump dates are smoothed with a Gaussian KDE and the mode is identified; the histogram summarizes these modes across all samples.

**Figure S8.**
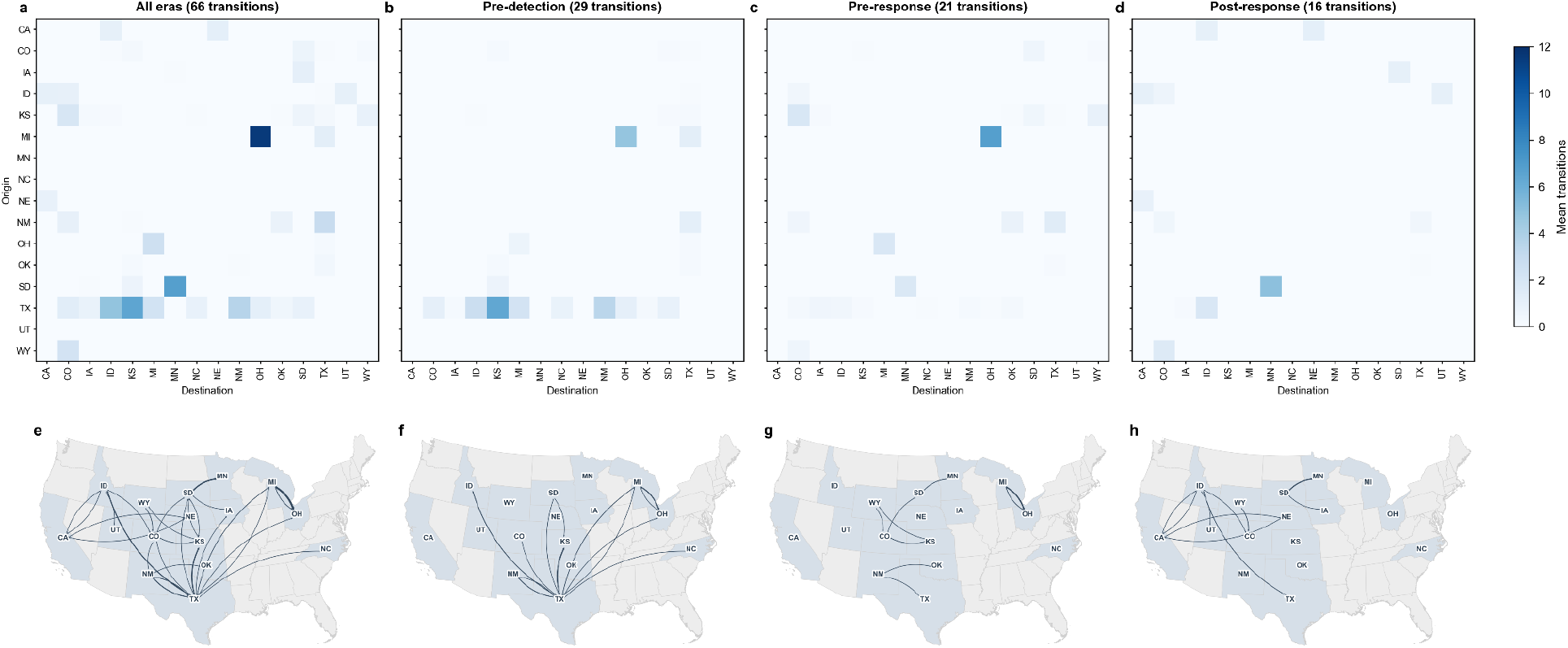
Phylogeographic estimates of H5N1 spread for each era. **a–d**, State-to-state transition matrices from the phylogeographic log-linear model. Each cell shows the posterior mean number of Markov jump transitions from origin state (row) to destination state (column). **e–h**, Maps of the contiguous US indicating transitions from state to state, with arrows scaled to the number of transitions. **a,e**, All transitions summed across the full study period (66 transitions). **b–d,f–h**, Transitions decomposed by era: (**b,f**) pre-detection (before 25 March 2024; 29 transitions), (**c,g**) pre-response (25 March 2024 – 29 April 2024; 21 transitions), and (**d,h**) post-response (after 29 April 2024; 16 transitions). Transition counts in each panel title are the posterior mean of per-sample totals. Color scale is shared across all heatmap panels; arrow width is scaled to the same maximum across all map panels. Texas as an origin dominates panels **b** and **f**, reflecting its role as the primary source of interstate spread during the pre-detection era. In panels **c** and **g**, Michigan emerges as the primary source during the pre-response era. Panels **d** and **h** show diffuse, lower-magnitude transmission with no single dominant source.

**Figure S9.**
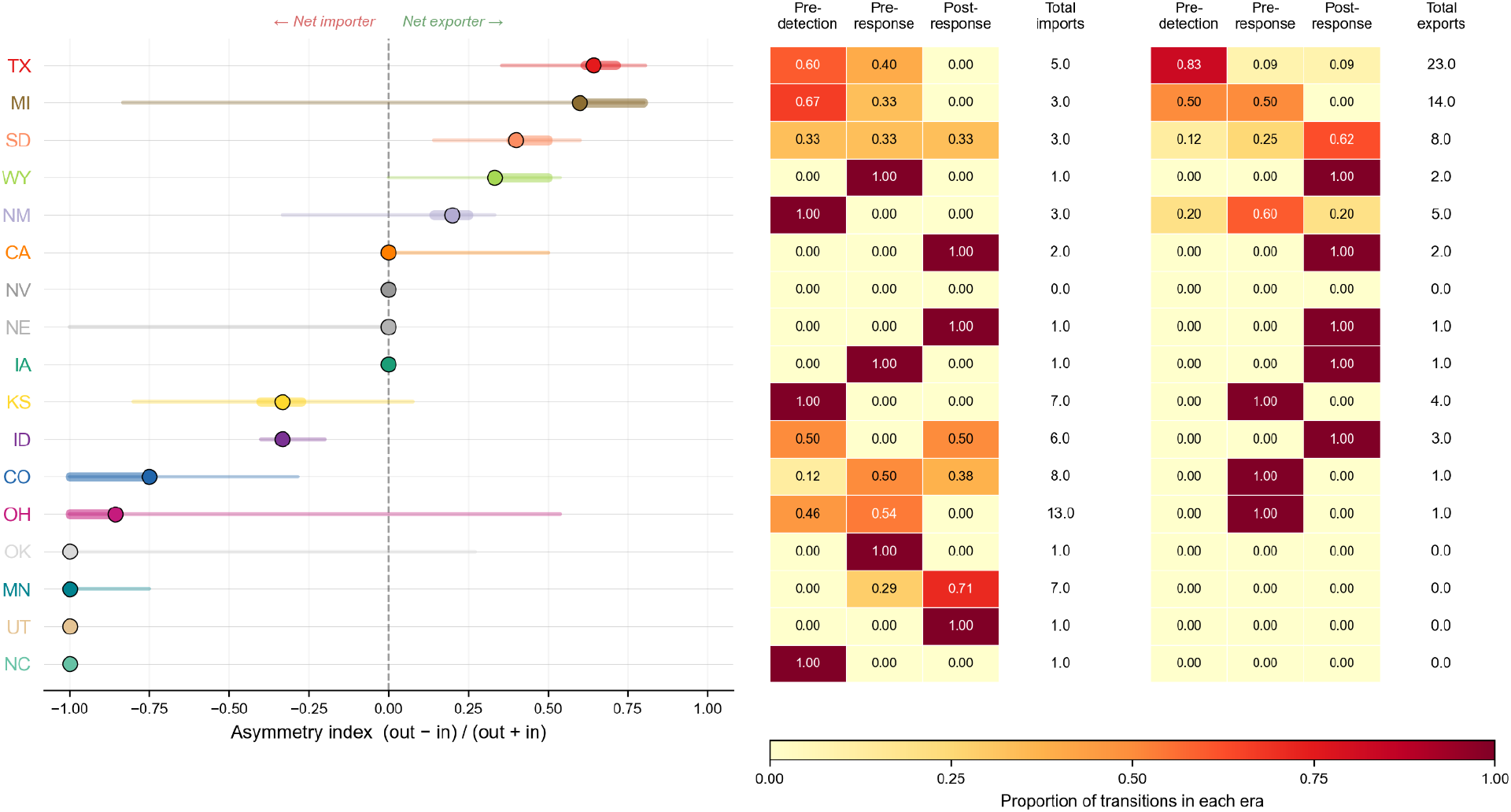
Transmission asymmetry index for 16 states. Left panel: for each posterior sample, the asymmetry index is computed as (out − in) / (out + in), where "out" and "in" are the numbers of Markov jump transitions originating from and arriving at each state, respectively. Values of +1 indicate a pure exporter (all transitions outward), −1 a pure importer (all transitions inward), and 0 balanced flow. States are sorted by posterior median asymmetry value; ties are broken by the upper 95% HPD bound. Dots show posterior medians; thick bars show 50% HPD intervals; thin bars show 95% HPD intervals. Middle and right panels: heatmaps showing the proportion of each state’s imports (middle) and exports (right) occurring in each era (pre-detection, pre-response, post-response), with the posterior median total number of imports or exports listed to the right of each heatmap. Texas is the strongest net exporter (median = 0.643, 95% HPD [0.355, 0.806]), with 83% of its exports occurring during the pre-detection period, consistent with its role as the phylogenetic root and primary source of early interstate spread. Michigan is the second-largest net exporter (median = 0.600), with exports split evenly between the pre-detection (50%) and pre-response (50%) eras; however, its wide 95% HPD [−0.833, 0.800] reflects substantial uncertainty, largely driven by the uncertain number of transitions between Michigan and Ohio. Minnesota, Utah, Oklahoma, and North Carolina are strong net importers (median = −1.0), reflecting states that received viral introductions but rarely seeded onward transmission.

**Figure S10.**
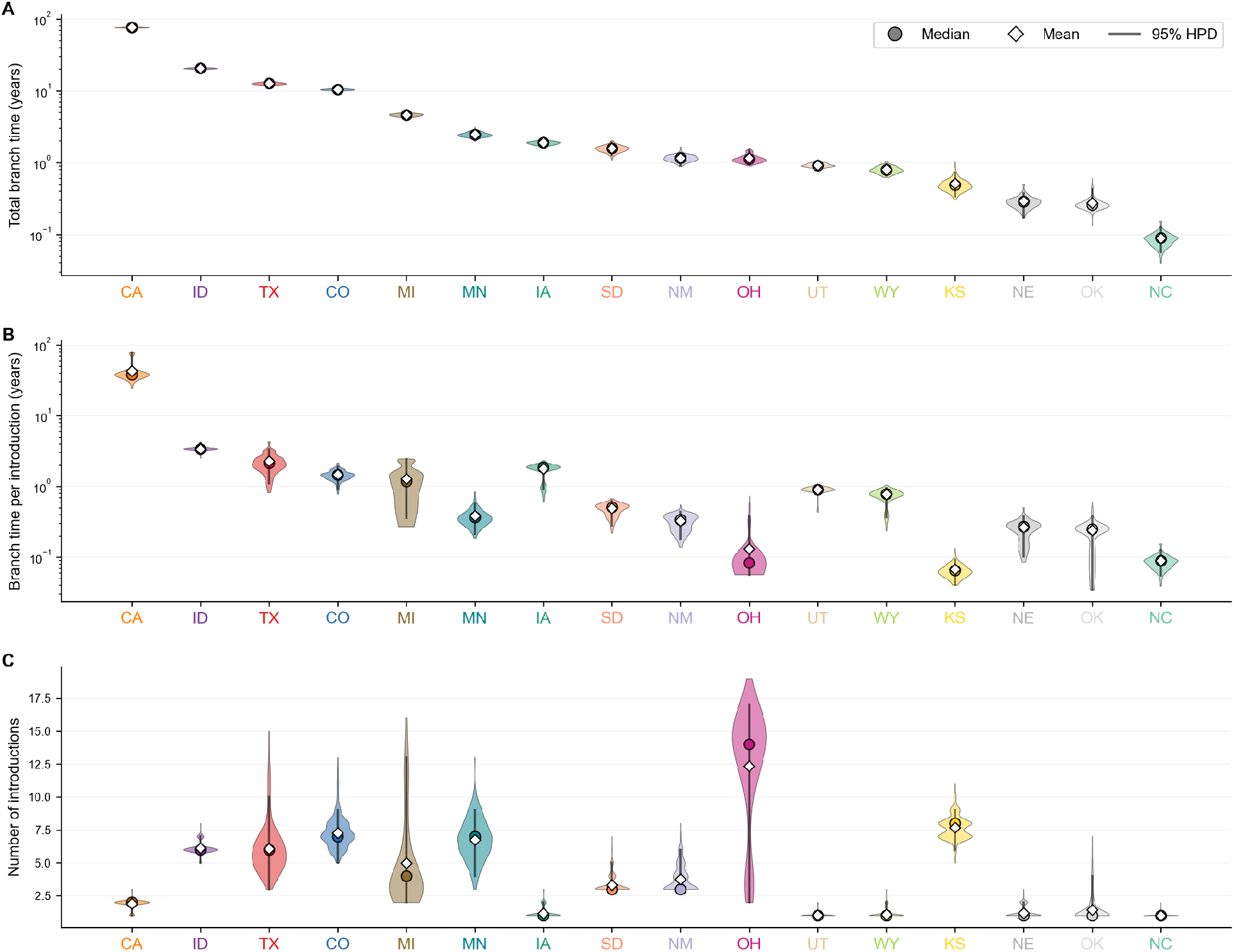
Phylogenetic persistence data. Posterior distributions of (**a**) total branch time for each of 16 states, ranked in descending order; (**b**) branch time per introduction for the same states, and (**c**) number of introductions into each state. The circle in each violin represents the median; the diamond represents the posterior mean; and the line represents the 95% HPD.

**Figure S11.**
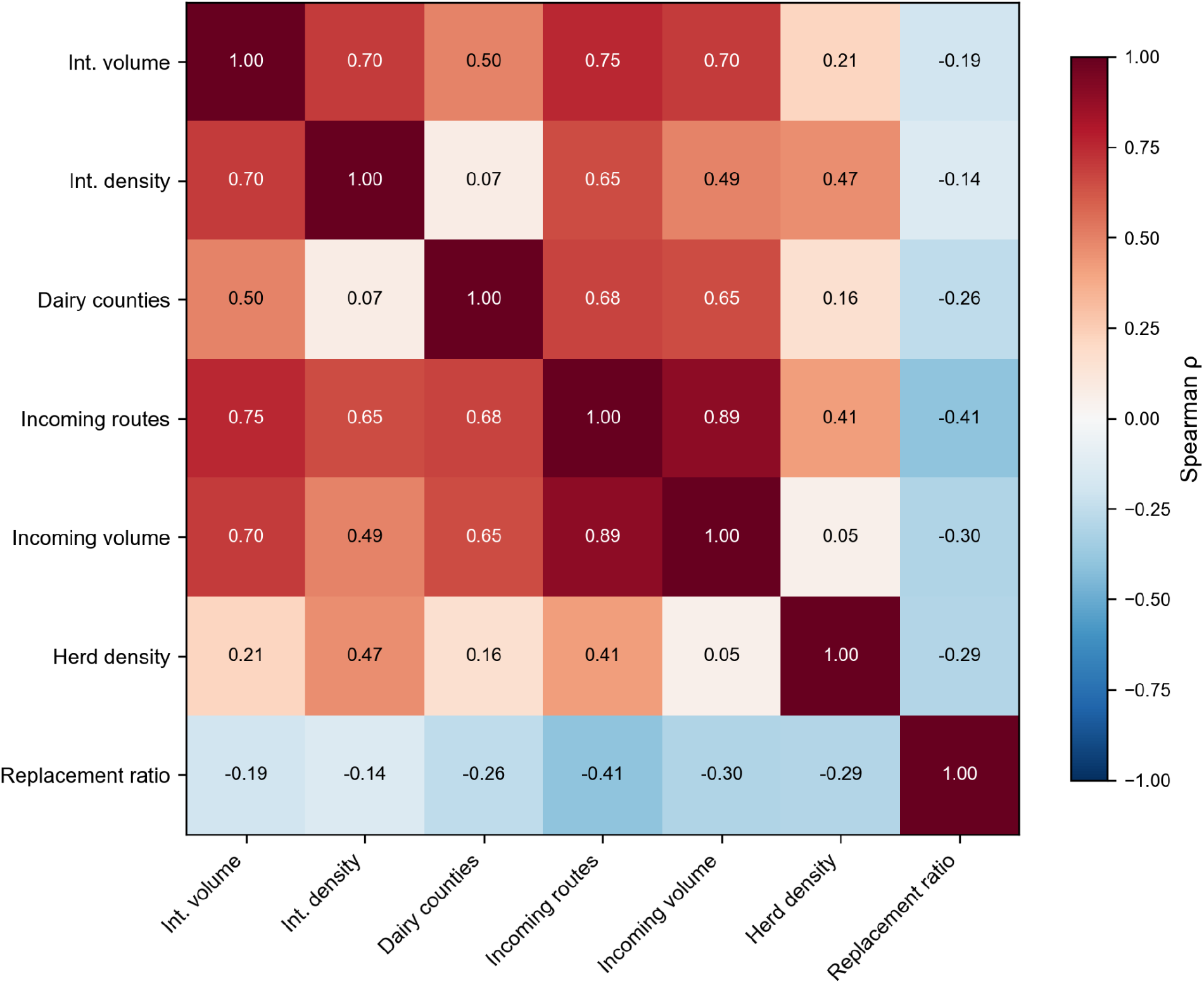
Spearman correlation matrix among the seven log-transformed network predictors. Internal volume and density are moderately correlated (ρ = 0.70), along with several other predictor pairs (e.g., internal density and incoming volume, ρ = 0.49). This, together with the small sample size (n=16) motivates the univariate analytical approach.

**Figure S12.**
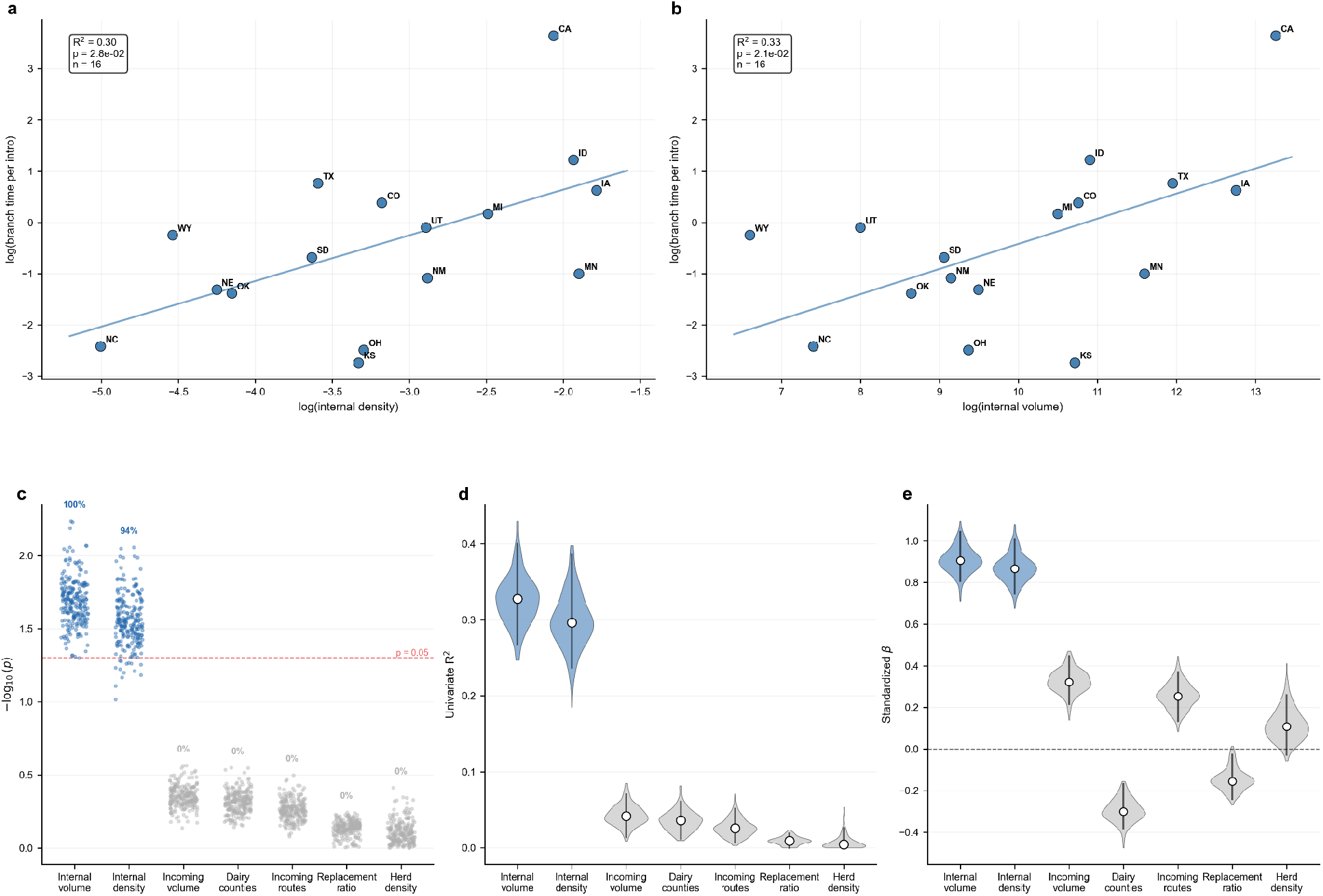
Persistence regression results using branch time per introduction as the response variable. **a**, Relationship between log(internal network density) and log(total branch time per introduction). **b**, Relationship between log(internal cattle movement volume) and log(total branch time per introduction). R² and P-values shown for regressions in **a–b**. **c**, Values of -log_10_(P) for each predictor across the posterior distribution; each point represents one posterior tree sample. Red dashed line indicates the significance threshold (P = 0.05), with points above the line denoting statistical significance. Percentages indicate the proportion of posterior trees with P < 0.05. **d**, Univariate R² for each of the seven network and dairy predictors (bars, 95% HPD). **e**, Standardized regression coefficients for each predictor. White dots indicate medians and thick bars indicate 95% HPDs. Blue violins denote predictors supported in ≥90% of posterior trees; grey violins denote predictors below this threshold. Dashed line indicates β = 0. Results are consistent with those obtained using total branch time as the response (Fig. 3), with internal cattle movement volume (median R² = 0.33, 100% of posterior trees significant) and internal network density (median R² = 0.30, 94% of posterior significant) remaining the dominant predictors, albeit with reduced effect sizes reflecting additional noise introduced by dividing by the posterior introduction count.

**Figure S13.**
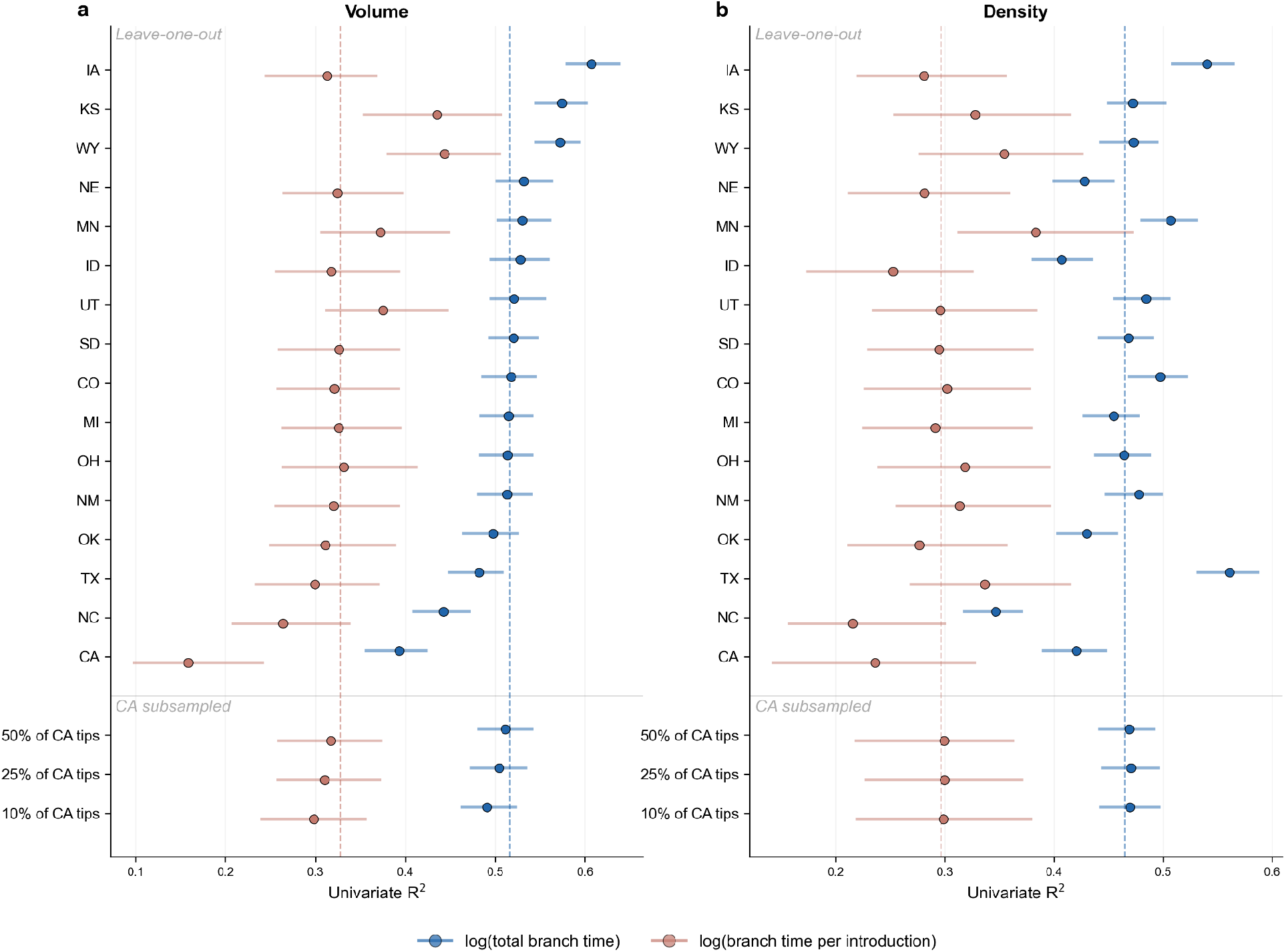
Leave-one-out robustness and California subsampling sensitivity of the volume and density persistence associations. **a**, Univariate R² for internal cattle movement volume when each state is sequentially excluded (n = 15). Dots show median R²; horizontal intervals span the 95% HPD. Dashed vertical lines indicate the full-data (n = 16) median R² for each response variable. Blue: log(total branch time); coral: log(branch time per introduction). Below the dividing line, California tips were randomly subsampled to 50%, 25%, and 10% of the original 2,232 tips, trees were pruned, and branch times and introduction counts were recomputed from the pruned trees before re-running the full 16-state regression. **b**, Same analysis for internal network density. Subsampling California to as few as 10% of its tips reduces the volume R² modestly while the density R² remains nearly the same, indicating that the associations are not driven by California’s disproportionate tip count.

**Figure S14.**
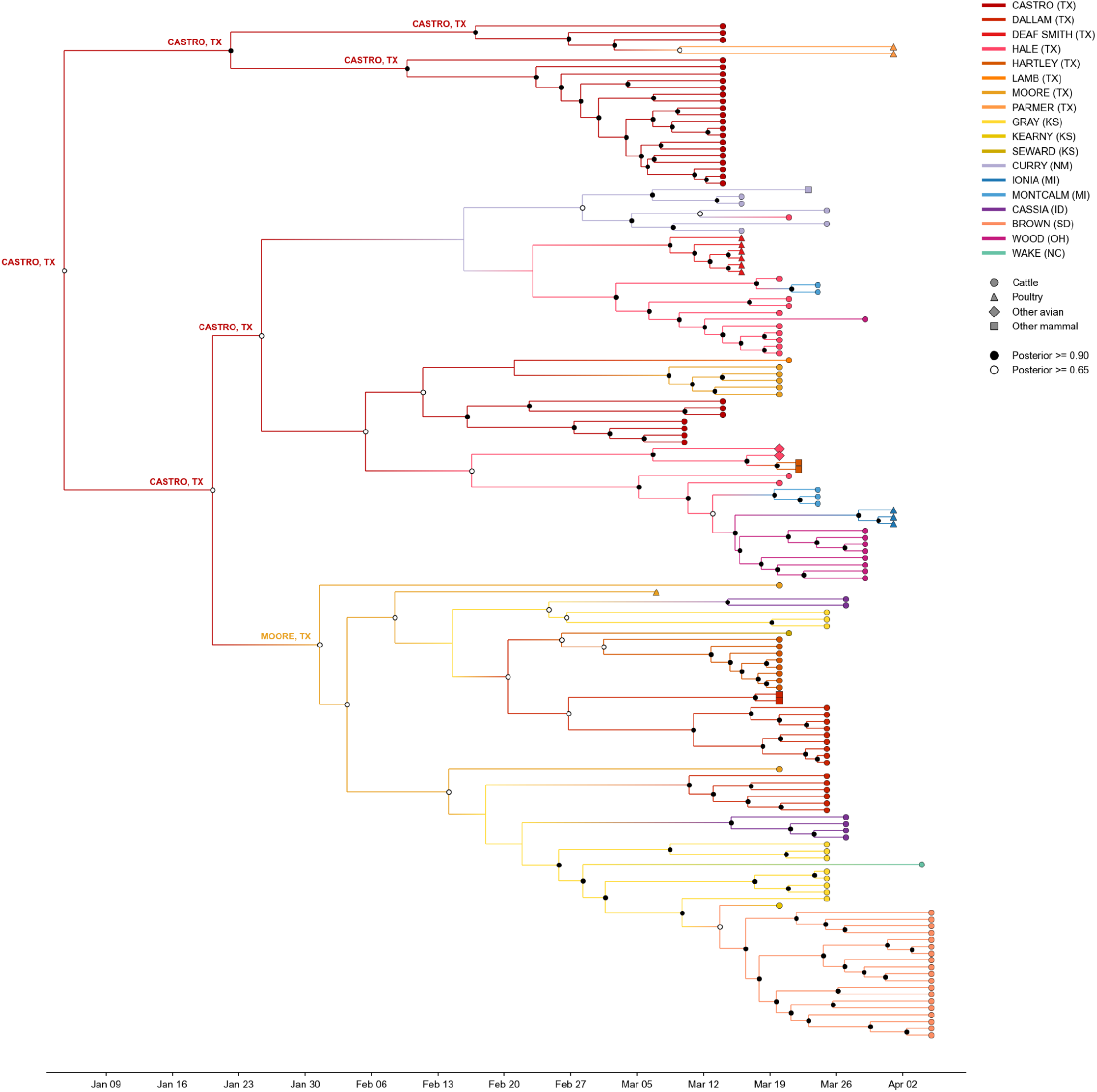
Time-scaled phylogenetic tree of 149 early outbreak virus genomes with county-level discrete trait reconstruction. Viruses sampled from non-cattle hosts were included because of how few genomes had county-level metadata, and the hosts can be considered sentinels for cattle transmission in those regions. Branches are coloured by the most probable county assignment. Circles at internal nodes indicate location posterior probability ≥0.65 (open circles) and ≥0.95 (filled circles). Tip shapes indicate host: circles (cattle), triangles (poultry), diamonds (other avian), squares (other mammals). The inferred location of the root, its two child nodes, and its four grandchild nodes are indicated.

**Figure S15.**
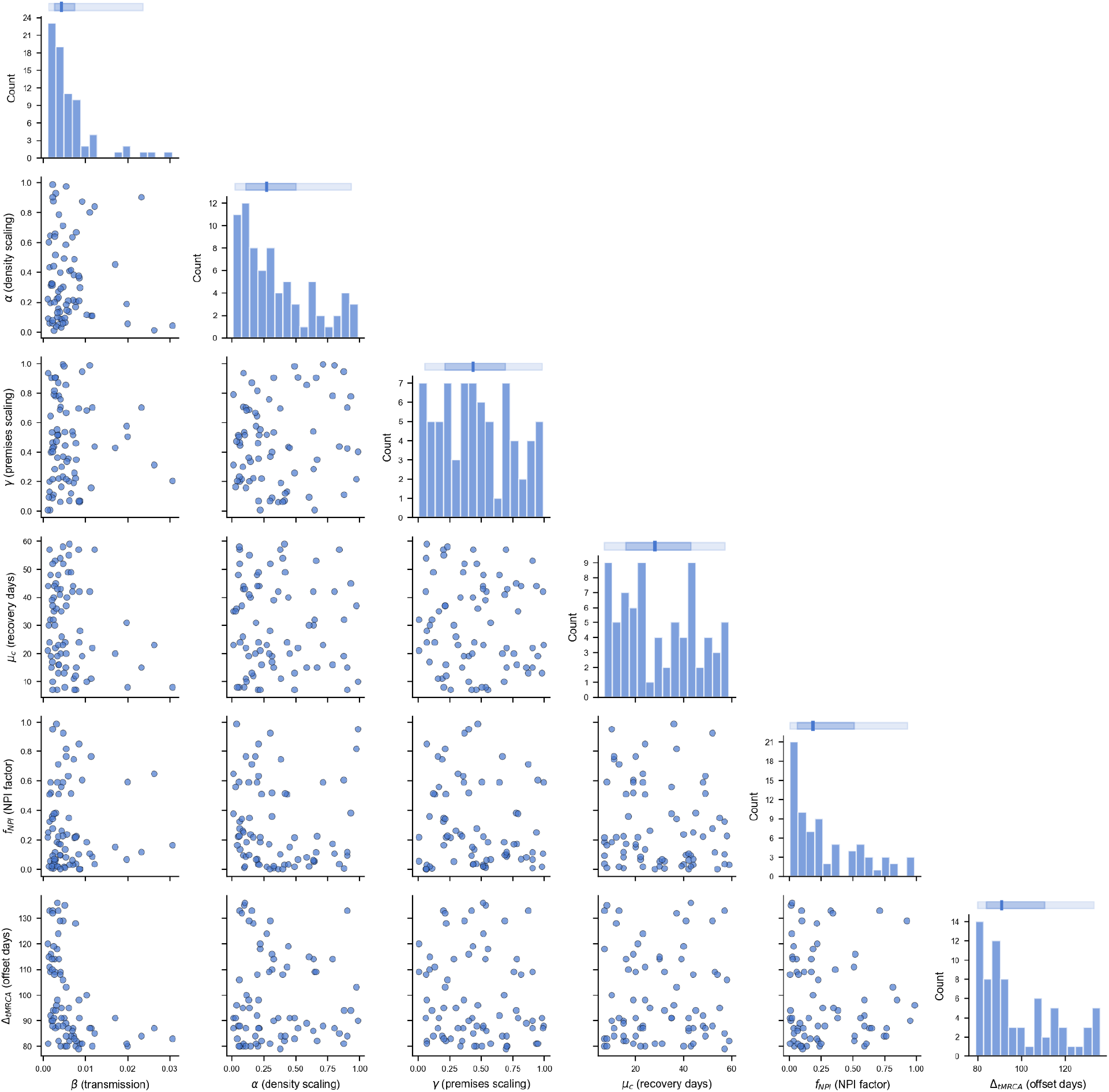
ABC-SMC posterior parameter scatter matrix. Pairwise relationships among the 75 accepted parameter sets from ABC-SMC calibration. Diagonal panels show marginal distributions for each parameter, with summary bars above (dark line, median; dark rectangle, interquartile range; light rectangle, 95% confidence interval). Lower triangle panels show pairwise scatter plots. The six calibrated parameters are: transmission rate (β), herd-size scaling of recovery (α), premises scaling of recovery (γ), base county recovery period in days (μ_c_), NPI reduction factor (f_NPI_), and tMRCA offset (days before the first Texas detection on 25 March 2024). The posterior spans a diverse range of parameter combinations, with β and tMRCA offset showing the most structured joint distribution (earlier start dates [greater tMRCA offsets] calibrate to lower transmission rates). The scaling and recovery parameters sample broadly across their prior ranges, consistent with being weakly constrained by the observed outbreak data.

**Figure S16.**
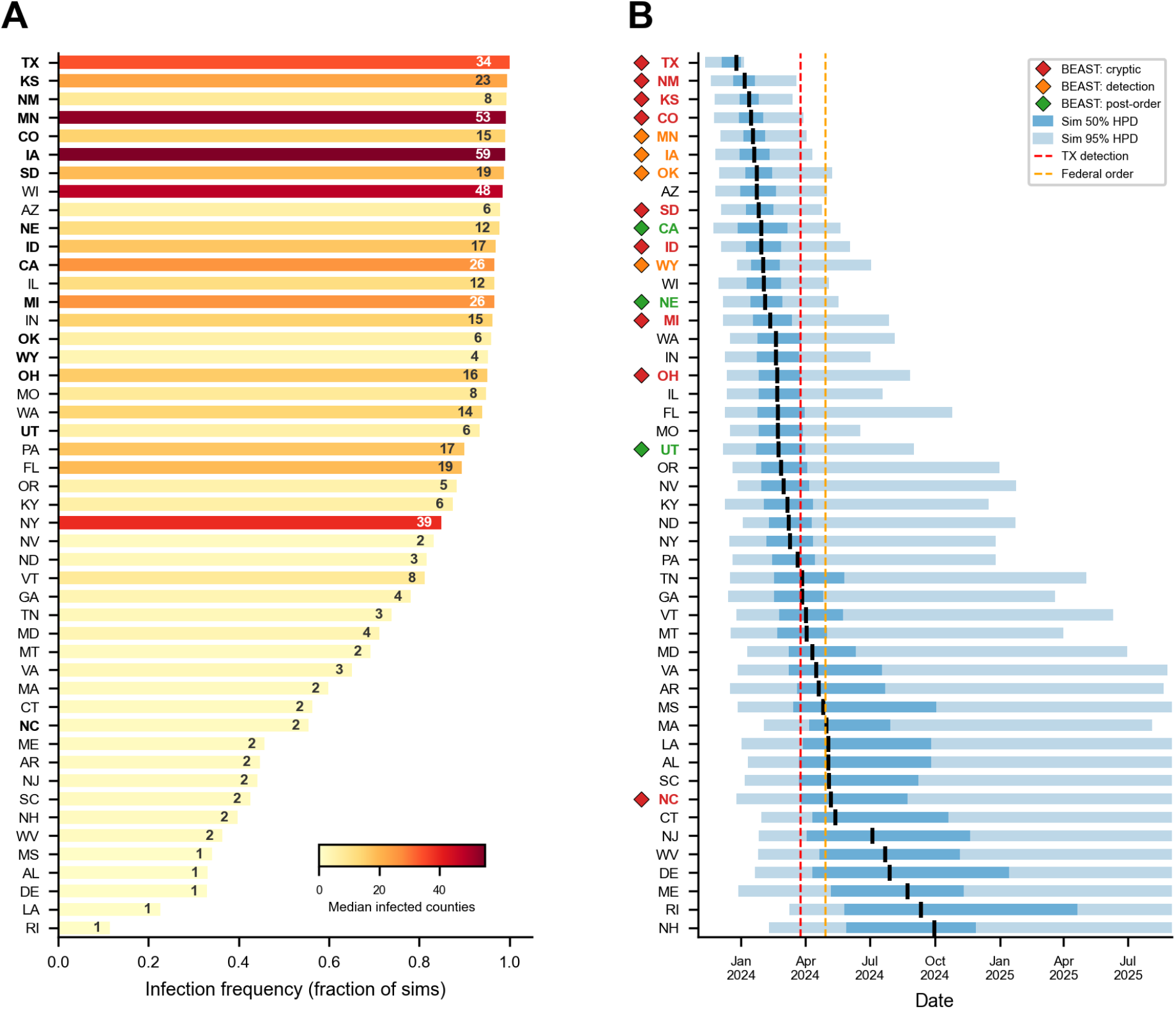
Simulation model validation. **a**, Infection frequency across 375,000 simulations incorporating non-pharmaceutical interventions, showing the fraction of simulations in which each state was infected. Bars are colored by the median number of infected counties (yellow–red gradient), with the median value indicated in each bar. Bold labels indicate the states included in the phylogeography. **b**, Simulated first-infection timing for each state, shown as 50% (dark blue) and 95% (light blue) highest posterior density intervals across simulations, with median indicated by a black tick. Colored diamonds mark the BEAST-estimated phylogeographic era for each state’s first introduction: pre-detection (red; before 25 March 2024), pre-response (orange; 25 March – 29 April 2024), post-response (green; after 29 April 2024). Vertical dashed lines indicate Texas detection (red) and the federal testing order (orange). The simulation model was calibrated via ABC-SMC using outbreak counts and detection timing.

**Figure S17.**
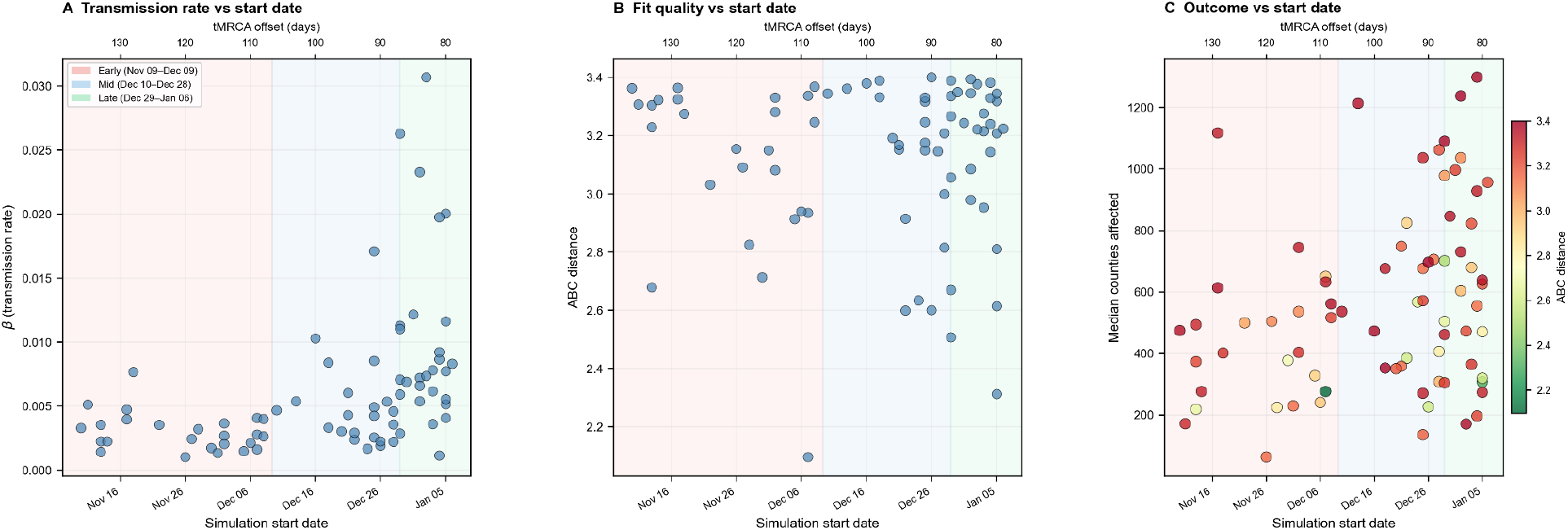
Start date sensitivity analysis. Relationship between simulation start date (tMRCA offset) and calibrated parameters, fit quality, and epidemic outcomes across 75 ABC-SMC accepted parameter sets. Start dates span 9 November 2023 to 6 January 2024 (79–137 days before the first Texas detection on 25 March 2024). Background shading indicates tertile groups: early (red; November 9–December 9), mid (blue; December 10–December 28), and late (green; December 29–January 6). **a**, Calibrated transmission rate (β) versus start date. Earlier start dates calibrate to lower β values, reflecting parameter compensation: longer cryptic periods require slower transmission to match observed data. **b**, ABC distance (goodness-of-fit) versus start date. Fit quality is stable across the full range of start dates, indicating that the ABC-SMC procedure successfully calibrates parameters regardless of assumed epidemic origin. **c**, Median counties affected (across 5,000 simulations per parameter set) versus start date, colored by ABC distance (green = better fit, red = worse fit). Previously inferred tMRCA for B3.13 in dairy cattle had an HPD of 9 November 2023–6 January 2024; the time of the parent of the MRCA (tPMRCA) had an inferred 95% HPD of 30 September 2023–22 November 2023. Our simulation start dates correspond to the tMRCA and overlap with the tPMRCA, and our results suggest that epidemic outcomes are broadly consistent across start dates, with no systematic dependence on assumed origin timing.

**Figure S18.**
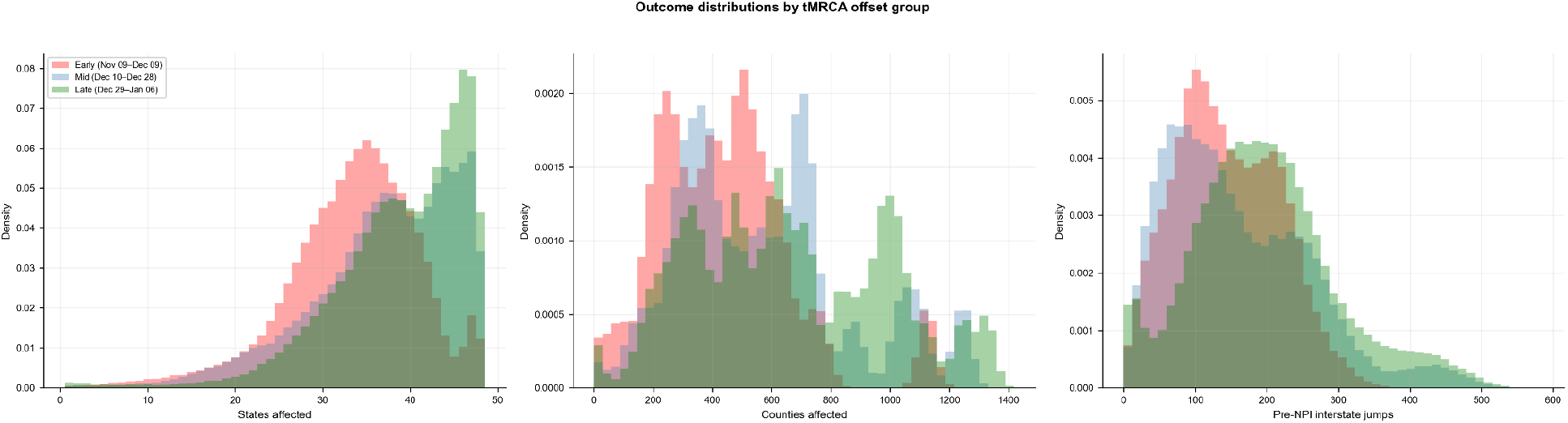
Stratified simulation outcome distributions by start date group. Density histograms of simulation outcomes stratified by tMRCA offset tertile: early (red; November 9–December 9, n = 125,000 simulations from 25 parameter sets), mid (blue; December 10–December 28, n = 110,000 from 22 parameter sets), and late (green; December 29–January 6, n = 140,000 from 28 parameter sets). Each parameter set was run on 1,000 networks × 5 replicate simulations under the actual NPI scenario (between-state movement restrictions imposed 35 days after the first Texas detection). a, Number of states affected. b, Number of counties affected. c, Number of pre-NPI interstate transmission events. Outcome distributions overlap substantially across all three groups, supporting the conclusion that simulation results are robust to the assumed start date within the calibrated prior range.

**Figure S19.**
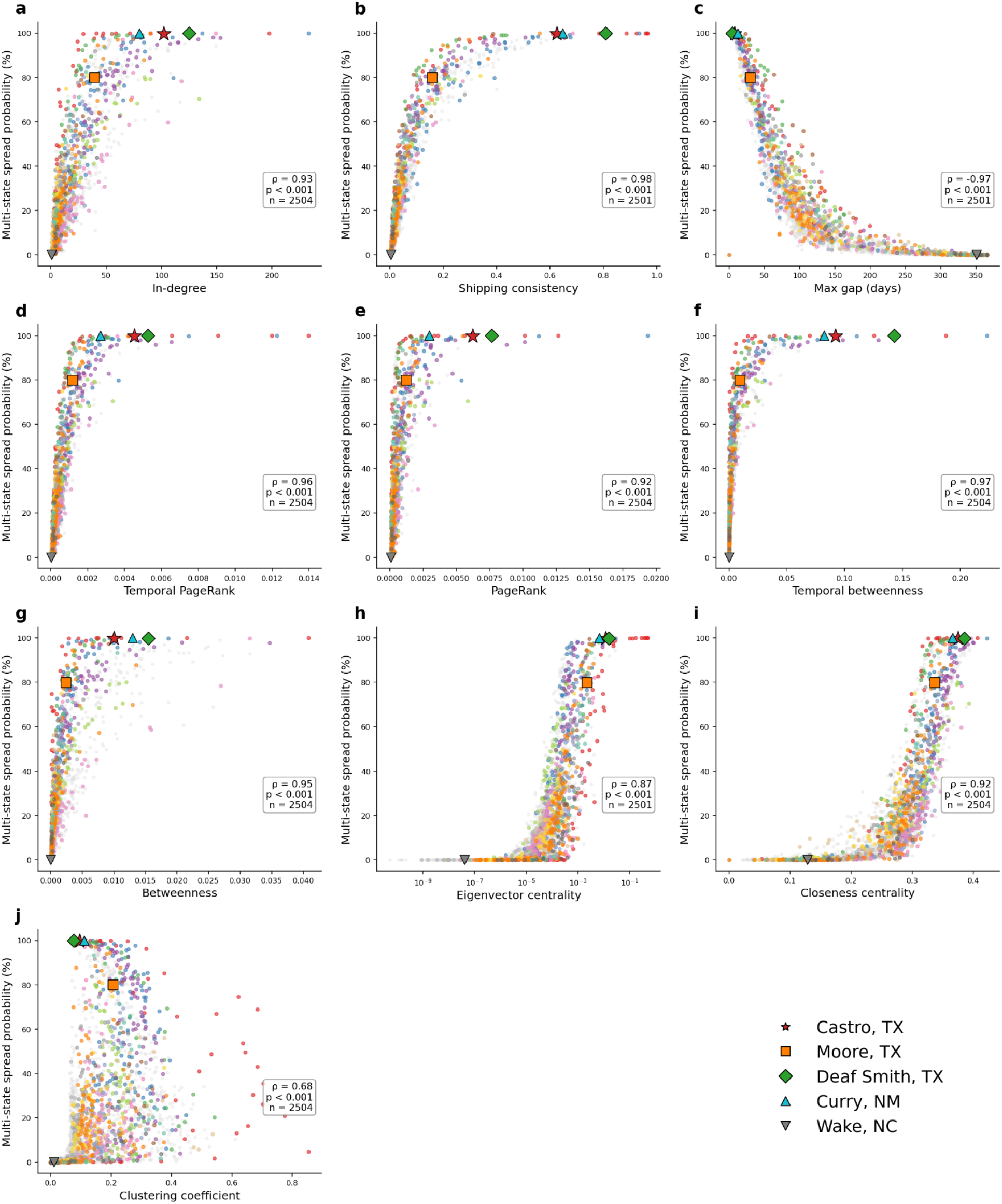
Network centrality metrics versus probability of multi-state spread. Each panel shows a different network metric (see Table S6 for definitions) plotted against the probability of multi-state spread for each of the seed counties across the 48 contiguous states, estimated from alternative origin simulations. Points are colored by state; highlighted counties are Castro (red star), Moore (orange square), and Deaf Smith (green diamond), Texas; Curry, New Mexico (light blue triangle); and Wake, North Carolina (gray inverted triangle). Spearman rank correlation (ρ) and sample size shown in each panel. **a**, In-degree. **b**, Shipping consistency. **c**, Maximum shipping gap. **d**, Temporal PageRank. **e**, PageRank. **f**, Temporal betweenness. **g**, Betweenness. **h**, Eigenvector centrality. **i**, Closeness centrality. **j**, Clustering coefficient. Eigenvector centrality (**h**) is displayed on a log scale; counties with eigenvector centrality of zero are excluded from that panel. Out-degree and temporal reachability are omitted here as they appear in Figure 4d and 4e, respectively. All metrics averaged across all 1,000 USAMMv3 dairy cattle network realizations. The majority of metrics were calculated using 2,504 counties with dairy cattle; three counties did not have outward shipments, resulting in a subset of metrics being calculated for 2,501 counties. Among the highlighted counties, Moore County had lower percentile ranks across all network features except clustering, reflecting a smaller set of more interconnected trading partners, while Castro, Deaf Smith, and Curry counties shipped to more destinations whose trading partners were less interconnected with one another. Wake County, from which there were likely no onward transmissions, expectedly approached the lowest percentile ranks across all metrics (also see Figure 4d–4f).

**Figure S20.**
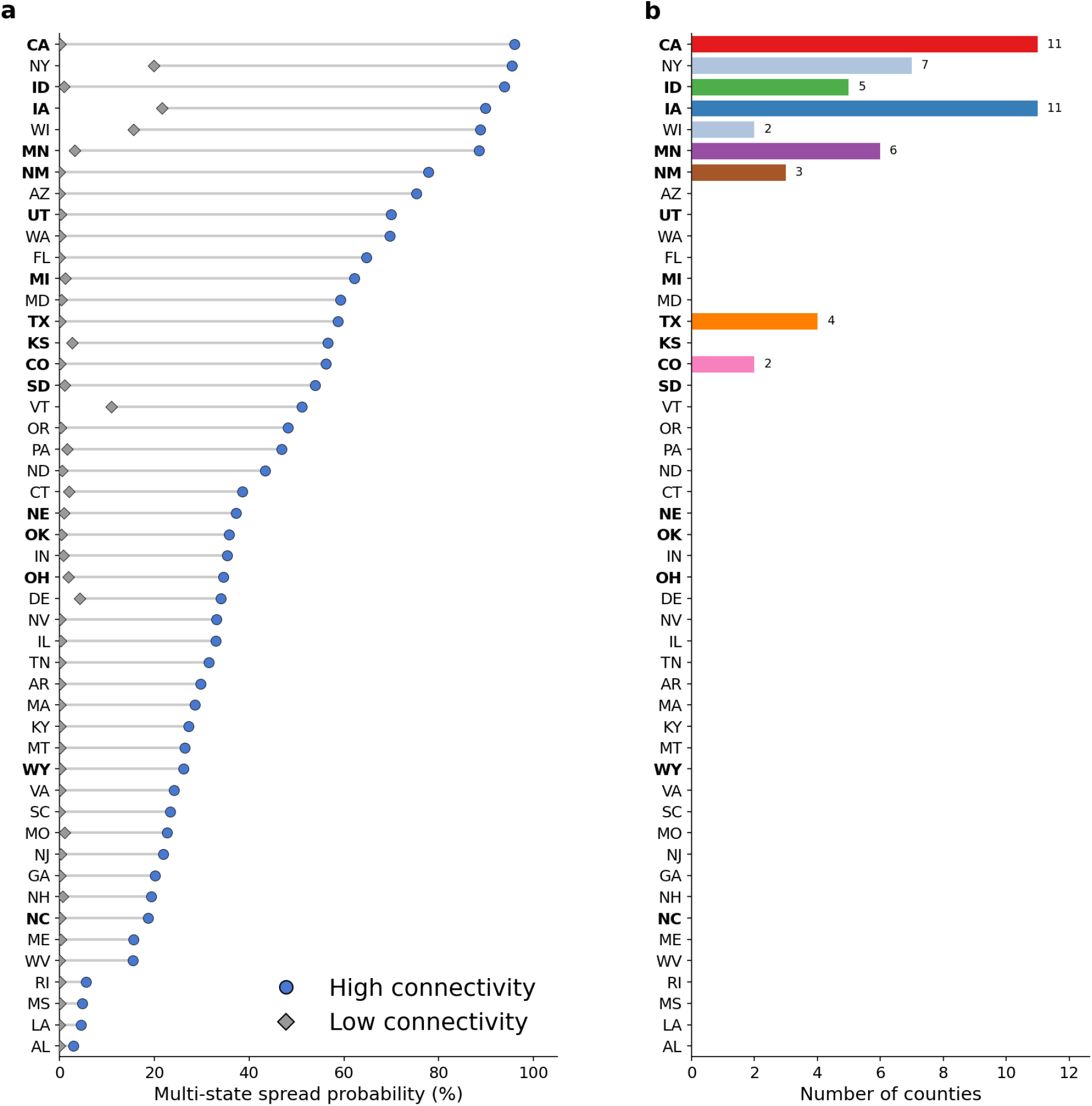
Multi-state spread probabilities at the state level. **a**, Barbell plot comparing average multi-state spread probabilities of low-connectivity and high-connectivity seed counties across the 48 contiguous states, with the 16 states included in the phylogeographic analysis in bold. For all states except Delaware (DE), high-connectivity counties are those with out-degree ≥75th percentile, whereas low-connectivity counties are those with out-degree ≤25th percentile. Since Delaware only has 3 counties with dairy cattle, we instead segregate into one high-connectivity county and one low-connectivity county. **b**, Barplot of the number of counties where multi-state spread probability (reaching ≥2 additional states) is ≥95%.

**Figure S21.**
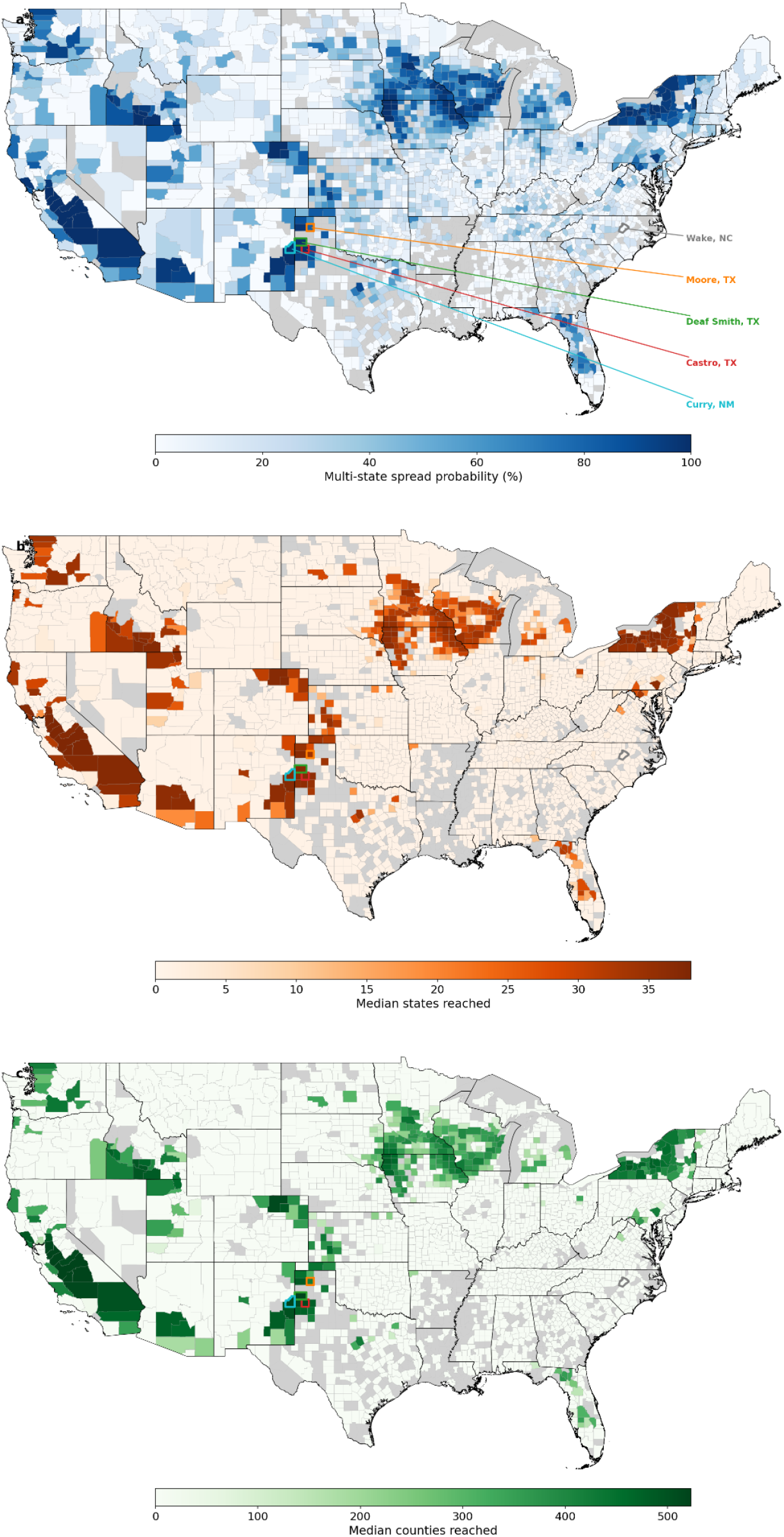
County-level multi-state spread probabilities. Map detailing probability of an outbreak reaching at least two other states (top), the median number of states reached (middle), and the median number of counties reached (bottom) for each county where an outbreak was initiated. Grey counties are non-dairy counties and were not included in the simulations. Castro County, Moore County, and Deaf Smith County in Texas, Curry County in New Mexico, and Wake County in North Carolina indicated.

**Figure S22.**
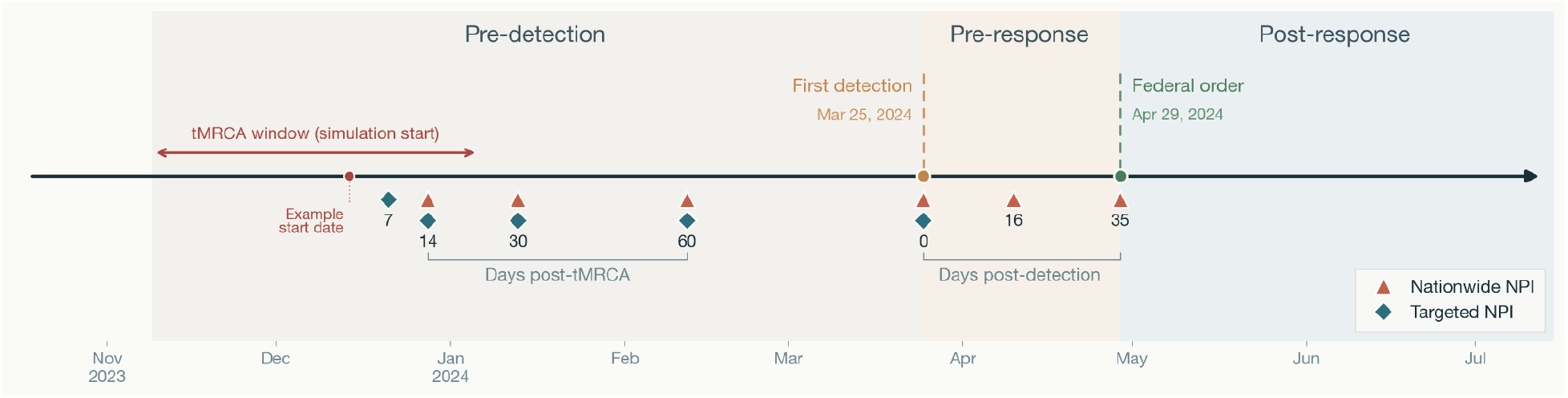
Schematic of counterfactual NPI scenarios. Each simulation was seeded with an infection in Castro County, Texas, at a time sampled from a previously inferred B3.13 tMRCA range (two-sided red arrow). The baseline scenario invoked a federal order restricting interstate cattle movement on the actual federal order date (29 April 2024, 35 days after the first confirmed detection), with effectiveness sampled from the ABC-SMC posterior (median 81.5% reduction; 50% CI: 48.7–93.5%, 95% CI: 6.9–99.7%). Nationwide NPI counterfactual scenarios varied the timing and strength of the nationwide NPI (Fig. S23): between-state strength scenarios applied 10–90% reductions in interstate transmission at the federal order date; timing scenarios applied a standardized 90% reduction at detection or 16 days post-detection; pre-detection surveillance scenarios assumed intervention at 7, 14, 30, or 60 days post-tMRCA with 90% reduction; and a combined within-and-between-state scenario combined 90% within-state and 90% between-state reductions at detection. Targeted NPI counterfactual scenarios focused on spatial scale and dairy cattle mobility (Fig. S24, Fig. S25), halting all transmission from targeted counties at either 14, 30, or 60 days post-tMRCA or at detection. Triangle markers indicate the timing of each nationwide NPI relative to the tMRCA and Texas detection dates. Diamond markers indicate the timing of each targeted NPI relative to the tMRCA and Texas detection dates.

**Figure S23.**
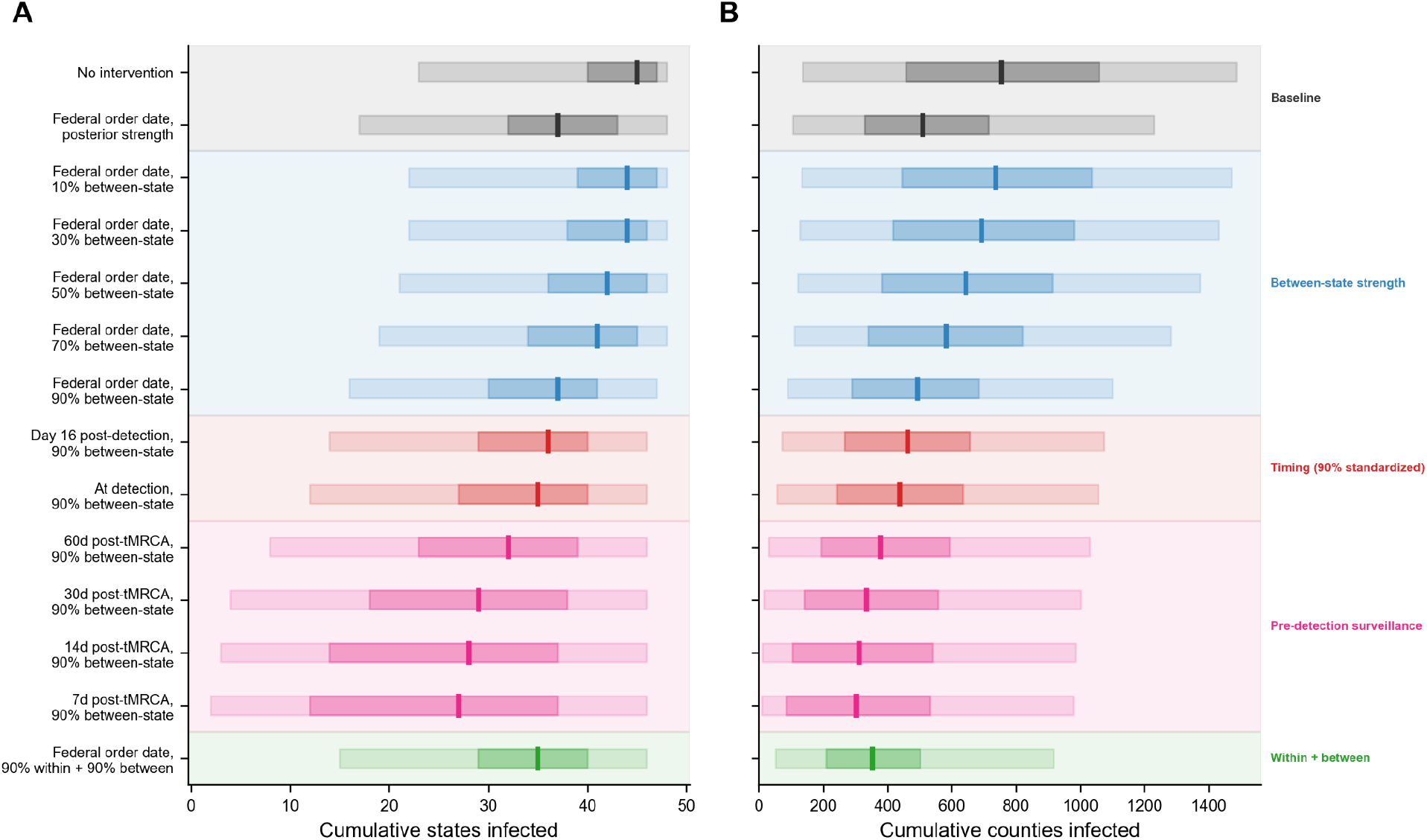
Non-pharmaceutical intervention scenario comparison. Cumulative states (**a**) and counties (**b**) infected after two years of simulated transmission across 14 counterfactual scenarios grouped into five categories. Vertical bars show medians; dark shaded regions show interquartile ranges (25th–75th percentile); light shaded regions show 95% confidence intervals (2.5th–97.5th percentile). Baseline scenarios include no intervention and the calibrated federal order (posterior effectiveness; see Figure S15 for the full distribution of the f_NPI_, which represents effectiveness in our simulations). Between-state strength scenarios vary the reduction in interstate transmission from 10% to 90%, all implemented at the federal order date. Timing scenarios vary the delay between detection and policy implementation at a standardized 90% between-state reduction. Pre-detection surveillance scenarios assume intervention at 7, 14, 30, or 60 days post-tMRCA with 90% between-state reduction. The within-and-between-state scenario combines 90% within-state and 90% between-state movement reductions at detection. See Table S7 for numerical values of the simulation outcomes.

**Figure S24.**
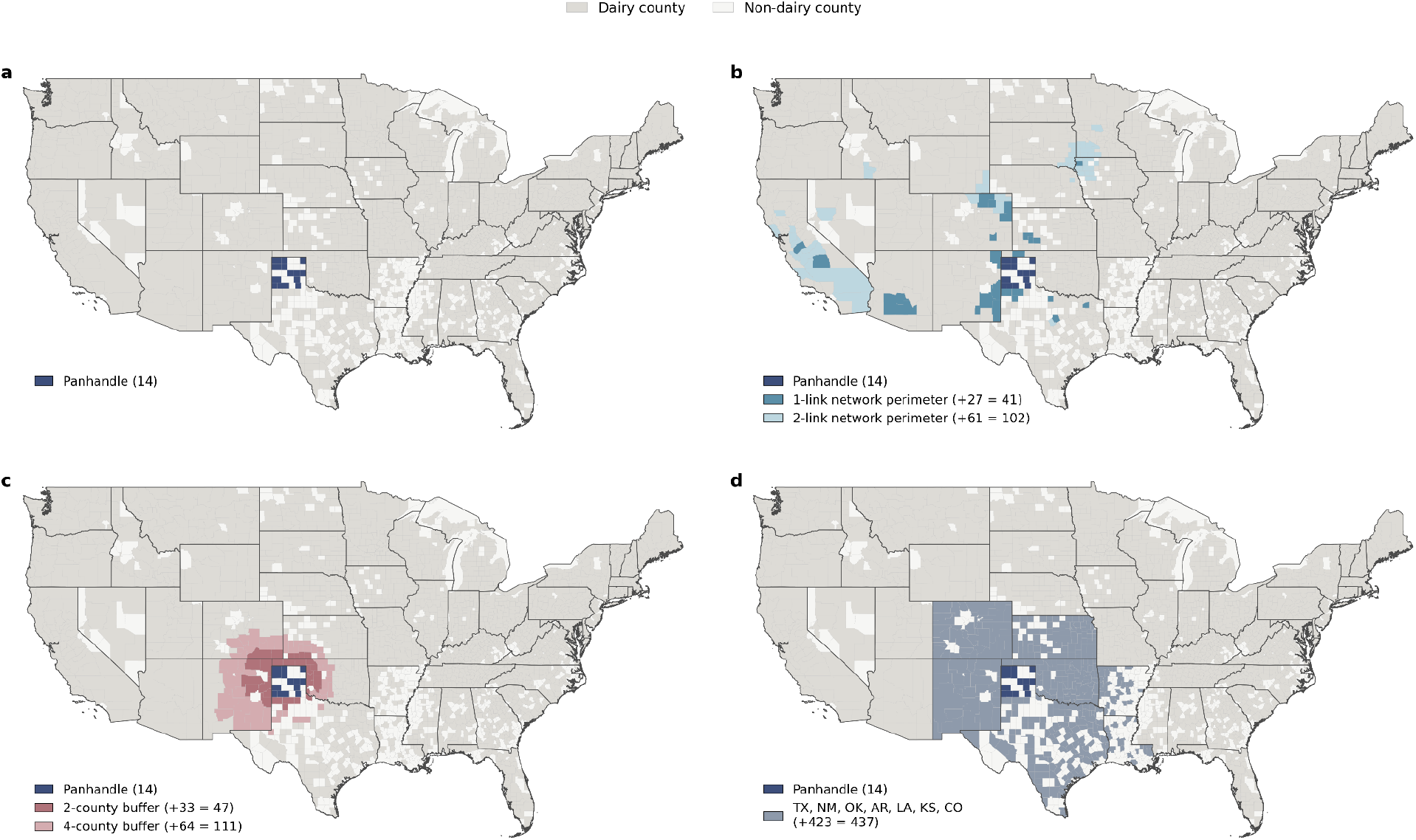
Counties targeted for rapid early intervention. **a**, The 14 counties with dairy cattle in the Texas Panhandle, targeted by an idealized (100% effective) intervention limiting between-county transmission. **b**, Counties targeted based on connectivity in USAMMv3 network realizations: direct shipping destinations of the Panhandle (1-link network perimeter) and counties reachable within two shipments (2-link network perimeter; see Methods). **c**, Counties targeted based on geographic proximity: a narrow perimeter of counties adjacent to the Panhandle dairy counties (2-county buffer) and a wide perimeter extending further outward (4-county buffer). **d**, Counties targeted based on regional proximity, including all dairy counties in Texas, Arkansas, Colorado, Kansas, Louisiana, New Mexico, and Oklahoma.

**Figure S25.**
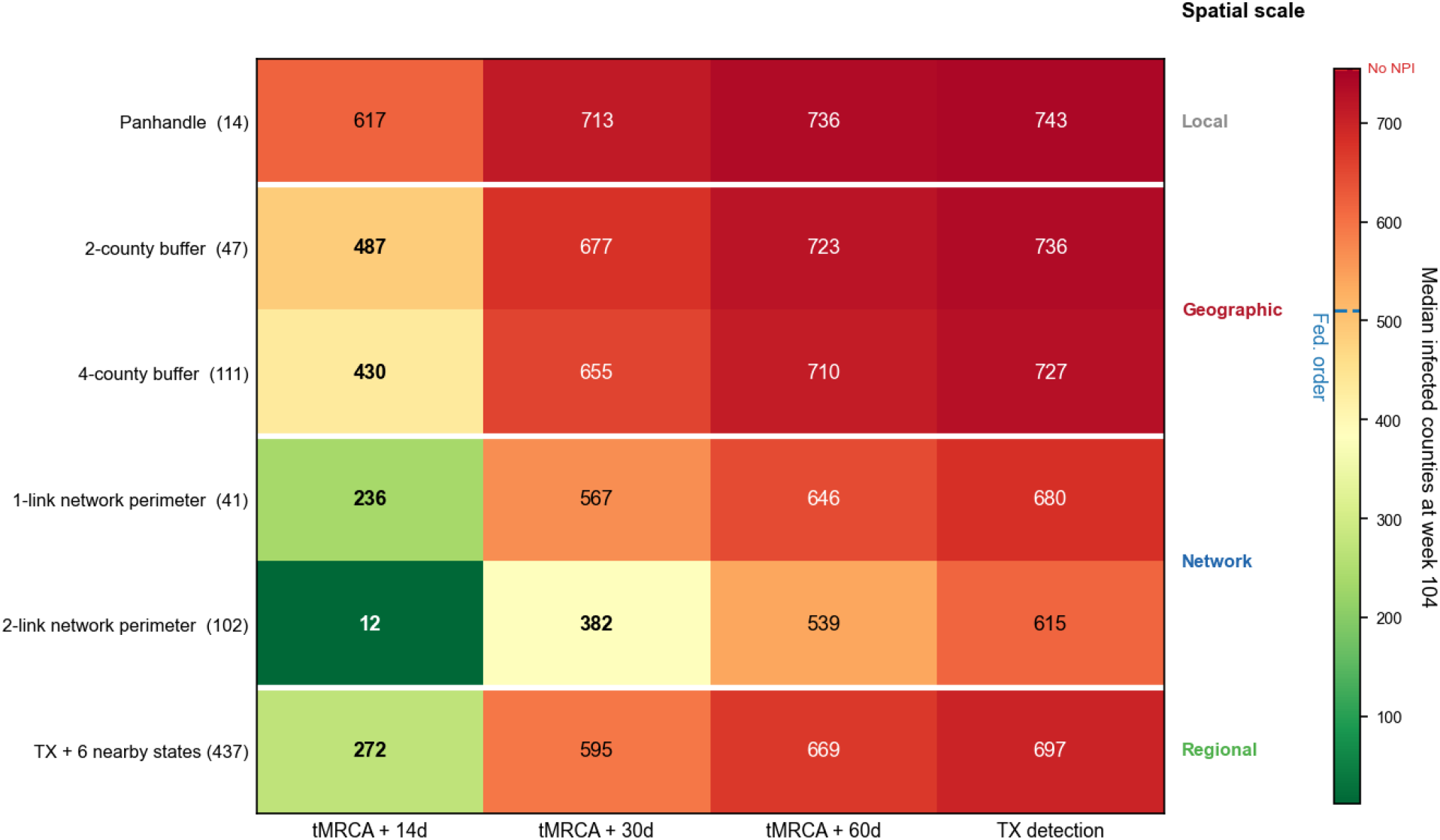
Targeted NPI comparison, focusing on the Texas panhandle. Cumulative counties infected after two years of simulated transmission across 24 counterfactual scenarios grouped into 4 categories. The interventions were 100% effective at limiting between-county transmission from the targeted counties. All groupings targeted counties in the Texas panhandle. The local grouping only targeted the 14 counties with dairy cattle in the Texas panhandle. The network grouping additionally targeted counties that were connected to those in the Texas panhandle via the USAMMv3 cattle movement network, whether they were directly connected by cattle trade (1-link network perimeter) or connected through one intermediary (2-link network perimeter). The geographic grouping targeted counties that formed either a narrow perimeter around the panhandle (2-county buffer) or a wide perimeter around the panhandle (4-county buffer). The regional grouping targeted all counties in Texas and nearby states. Number of counties targeted indicated in parentheses. Columns indicate whether the policy was implemented 14 days post-tMRCA, 30 days post-tMRCA, 60 days post-tMRCA, or at time of detection of the first Texas outbreak (102 days post-tMRCA). The colour scale is capped at 755, the median number of counties infected under the counterfactual with no NPI implemented. The blue dashed line indicates the median number of counties infected when the federal order was implemented. Refer to Figure S24 for maps indicating the counties targeted for each of these strategies. See Table S7 for additional numerical values of the simulation outcomes.

